# Decoding the Structural and Functional Impact of the Leukaemia-Associated A338V Mutation in GPR183

**DOI:** 10.64898/2026.03.30.715362

**Authors:** Louise Andersson, Patryk A. Wesolowski, Laura Jahrstorfer, Alexander De Rosa, Tomas Heger, Vilmos Neuman, Adam K. Sieradzan, David J. Wales, Paweł Kozielewicz

## Abstract

G protein–coupled receptors rely on dynamic conformational changes to coordinate G protein activation and recruitment of regulatory transducers such as G protein–coupled receptor kinases and β-arrestins. The chemotactic receptor GPR183 has been implicated in a context-dependent role in hematological malignancies. Here, we investigated the impact of A338V mutation located within the C-terminal tail of GPR183. This mutation is associated with acute myeloid leukaemia. Using bioluminescence resonance energy transfer-based assays in HEK293A cells, we assessed receptor-proximal signaling events. The A338V variant displayed preserved agonist potency and comparable agonist-induced Gi activation relative to wild type, although constitutive activity towards Gi was modestly reduced. In contrast, recruitment of GRK2 and β-arrestin2 was consistently impaired across multiple assay configurations. These differences were not attributable to altered receptor abundance, as the C-tail untagged mutant exhibited increased plasma membrane expression despite reduced regulatory transducer engagement. While intramolecular conformational biosensor measurements revealed subtle differences in global receptor conformation between WT and A338V, extensive molecular dynamics simulations supported the altered conformational sampling of the C-terminal tail in the A338V variant. Together, these data support a model in which the A338V substitution selectively alters C-terminal structural dynamics, impairing GRK2 and β-arrestin2 recruitment while preserving G protein activation.

## Introduction

G protein-coupled receptors (GPCRs) constitute the largest family of membrane proteins involved in signal transduction and represent major pharmacological targets [1]. Their functional importance reflects their central role in translating extracellular stimuli into intracellular responses through conserved conformational rearrangements within the seven-transmembrane helical bundle [2, 3]. GPR183 (EBI2) is a Class A orphan GPCR that regulates immune cell migration in response to oxysterol gradients [4, 5]. GPR183-mediated migration depends on both Gi activation and β-arrestin recruitment [6]. Notably, the functional consequences of GPR183 expression and activation in pathophysiological settings are context-dependent: the receptor has been associated with beneficial roles in diffuse large B-cell lymphoma (DLBCL) [7], yet it appears to exert detrimental effects in acute myeloid leukemia (AML) [8]. Along these lines, a missense mutation - substitution of alanine with valine - at position 338 (A338V) in the C-terminus end (C-tail) of GPR183 has been identified as a rare variant specifically in AML through genomic sequencing efforts [9-11]. This substitution introduces a bulkier, aliphatic branched side chain that may influence local conformational flexibility or alter interaction interfaces. GPCR C-terminal domains are often conformationally dynamic yet critically involved in receptor desensitization, trafficking, and β-arrestin-mediated signaling. [6, 12-14].

Advances in structural biology have illuminated the molecular mechanisms of GPCRs by resolving high-resolution structures of many receptors bound to diverse ligands and signaling partners [15]. The advent of AlphaFold has further revolutionized the field by enabling remarkably accurate structural predictions across nearly all GPCRs [16]. Yet, GPCRs function as dynamic ensembles rather than static entities. Their conformational landscapes underpin key processes such as ligand selectivity and the adoption of active states necessary for recognition and coupling to heterotrimeric G proteins, β-arrestins, and GPCR kinases (GRKs). Each state is defined by distinct lifetimes and interconversion probabilities, modulated by cellular and environmental contexts. Capturing GPCR functional complexity, including transducer specificity, biased signaling, and allosteric regulation, thus demands integrating static structural snapshots with insights into receptor and transducer dynamics [17].

Intrinsically disordered regions (IDRs), notably the receptoŕs loops and C-tail, further fine-tune GPCR signalling [18]. IDRs are highly flexible and heterogeneous, often truncated or unresolved in X-ray or cryo-EM structures due to insufficient electron density [19]. AlphaFold struggles to capture the full conformational ensemble of such dynamic segments. Here, the GPR183 A338V mutation occurs within this unstructured/dynamic C-tail region. The structural and regulatory consequences of the A338V mutation remain undefined so far. Given the sensitivity of GPCR regulatory mechanisms to structural perturbation, alterations within the C-tail may affect receptor stability or protein–protein interactions, as already shown with several C-tail deletions in GPR183 [6]. Here, we hypothesize that the A338V substitution alters the conformational dynamics of the GPR183 C-terminal tail, leading to changes in proximal signaling events, including G protein activation and/or β-arrestin recruitment, compared to the wild type (WT) receptor. Complementary wet-lab experiments, including β-arrestin and GRK recruitment and G protein activation, and molecular dynamics simulations, were used to evaluate the mutation’s impact on receptor structure and function. To address this, we performed bioluminescence resonance energy transfer (BRET)-assays and combined them with AlphaFold-derived models of unresolved IDRs, including the C-tail region harboring the A338V mutation, which underwent molecular dynamics simulations in explicit lipid bilayers and solvent to probe their structural dynamics and potential functional effects.

As such, the aim of this study is to investigate the structural impact of the A338V substitution within the C-tail of GPR183 by comparing it to WT, with the goal of improving our understanding of how this leukaemia-associated variant may influence receptor function

## Material and methods

### Structural Preparation and Model Reconstruction

Cryo-EM structures of human GPR183 [20] contain substantial unresolved regions that preclude direct use in molecular simulations. The inactive structure (PDB ID: 7TUY) lacks residues 1-23, 94-99, 175-179, 187-192, 226-235, 275-280, and 321-361 (C-tail), while the active structure (PDB ID: 7TUZ) lacks residues 1-21, 94-99, 174-178, 317-361 (C-tail). These missing segments include intracellular loops and large portions of the C-terminal region. Notably, the leukaemia-associated A338V mutation is located within the unresolved C-terminus, necessitating full-length model reconstruction. The inactive structure (7TUY) contains a BRIL insertion replacing ICL3, along with anti-BRIL and anti-Fab nanobody domains used for cryo-EM stabilisation. All non-native domains were removed, the native ICL3 sequence restored, and the bound inverse agonist (PDB ID: KKF) deleted. For the active structure (7TUZ), the heterotrimeric. G protein and activating oxysterol ligand were removed to ensure consistency across simulations.

### Loop Reconstruction and Mutation Modelling

The experimental structures used for active and inactive states contained sequence differences (e.g., replaced sequence in ICL3, residues 226-235), resulting in different sequences being used for AlphaFold3 [21] input. The resulting structures were subsequently unified and refined using Modeller’s functionality ModLoop [22].Rest of the missing residues were reconstructed using comparative modelling. Predictions from AlphaFold2-Multimer [23], AlphaFold3 [21], and MODELLER [22, 24] were evaluated based on stereochemical quality and compatibility with resolved coordinates. Final models (AlphaFold3) were selected according to structural consistency and absence of steric clashes. The A338V substitution was introduced into both active and inactive conformations followed by local refinement. For the luciferase-attached model, we generated structures using AlphaFold3 with the full amino-acid sequence of GPR183 fused to Nluc, and used the resulting model for subsequent simulations. Six systems were constructed: GPR183 WT inactive, GPR183 WT active, GPR183-Nluc WT, GPR183 A338V inactive, GPR183 A338V active, and GPR183-Nluc A338V.

### Molecular dynamics

All systems were embedded in POPC bilayers using the Packmol–Memgen protocol [25], solvated with explicit TIP3P water [26] and neutralised to 0.15 M NaCl. A minimum 10 Å buffer was maintained between the protein and periodic box boundaries. Proteins were described with the AMBER ff14SB force field [27], lipids with Lipid17 [28] and water with the TIP3P model [26]. Molecular dynamics simulations followed previously established protocol [29]. Each system underwent two-stage energy minimisation: (i) 1000 steps (400 steepest descent, 600 conjugate gradient) with harmonic restraints (k = 100 kcal mol⁻¹ Å⁻²) applied to all solute atoms, followed by (ii) 2000 steps (800 steepest descent, 1200 conjugate gradient) with identical restraints applied to heavy atoms only. Equilibration was performed in three stages: (i) heating from 10 to 300 K over 10 000 steps (1 fs, NVT ensemble) with restraints on heavy atoms; (ii) 100 000 steps (1 fs, NPT ensemble) to stabilise density, with heavy-atom restraints maintained; and (iii) 400 000 steps (2 fs, NPT ensemble) with restraints applied to Cα and Cβ atoms. Production simulations were conducted in the NVT ensemble at 300 K using a Langevin thermostat, without restraints. Three independent trajectories of 100 × 10⁶ steps (2 fs timestep) were generated for each system, corresponding to 600 ns total sampling per system. In all explicit solvent simulations, the Particle Mesh Ewald method with a cutoff for nonbonded interactions set at 9 Å was used to speed up calculations.

### Molecular dynamics analysis

Trajectory analysis was performed to characterise residue interactions, conformational heterogeneity, and the underlying energy landscape. Conformational clustering was carried out using the DBSCAN algorithm as implemented in cpptraj [30], yielding representative structures corresponding to the most highly populated states. Secondary structure evolution along the trajectories was analysed using the DSSP algorithm in cpptaj.

Energy landscape organisation was examined using the MDDG framework [31] in combination with disconnectivity graph analysis [32, 33]. MDDG provides a coarse-grained representation of the sampled energy landscape by constructing disconnectivity graphs from the potential energy time series. Proxy minima and transition states were identified via Savitzky–Golay smoothing of the energy signal [34]. In disconnectivity graphs, energy increases along the vertical axis, while branch termini correspond to local minima. Minima are arranged horizontally to reflect funnel organisation, with superbasins defined by connectivity below a specified energy threshold [32, 33].

Solvent-accessible surface area (SASA) was computed to quantify global conformational changes using cpptraj, alongside hydrogen bond analysis. For each system, the SASA of the C-tail of GPR183 was calculated from three independent 200 ns simulations, yielding a total of 6000 frames. Statistical uncertainties were estimated via block averaging with a block size of 200 frames, exceeding the correlation time of the C-tail SASA, ensuring that each block represents an effectively independent sample and providing reliable standard errors of the mean. The same procedure was applied to assess the significance of disorder differences between systems.

### Cell culture

HEK293A cells and RAW264.7 cells were kept in an incubator in T-75 flasks with 10 ml DMEM, containing 10% FBS (Sigma) and 1% PenStrep, (Sigma) which was changed every 2-4 days. The absence of mycoplasma contamination was routinely confirmed by MycoStrip’s (InvivoGen) patented isothermal PCR detecting 16S ribosomal RNA of mycoplasma in the media after 2–3 days of cell exposure

### DNA constructs, cloning and mutagenesis

HiBiT-GPR183 plasmid DNA was generated with Gibson cloning using a codon-optimized GPR183 from GPR183-Tango plasmid DNA (#66342 Addgene, deposited by Bryan Roth) as an insert and HiBiT-FZD6 with a 5-HT3A signal peptide plasmid DNA as a backbone [35]. In the newly generated construct, the GPR183 insert sequence replaced FZD6 insert sequence. In this construct, the N-terminally cloned HiBiT tag (GTGAGCGGCTGGCGGCTGTTCAAGAAGATTAGC) is followed by a GS linker (GGATCC, BamHI site). HiBiT-GPR183-Nluc construct was generated using Gibson cloning, inserting Nluc from Nluc-FZD6 [36] onto the C-terminus of HiBiT-GPR183, without a linker. HiBiT-GPR183-mNG-Nluc plasmid DNA constructs were generated using Gibson cloning, inserting mNeonGreen (mNG) from mNeonGreen-APEX plasmid DNA (#202591 Addgene, deposited by Reuben Harris) at T233 position in HiBiT-GPR183-Nluc. Plasmid DNA constructs encoding the GPR183 A338V and Nluc R164Q mutant were generated with a GeneArt Site Directed mutagenesis kit (Thermo Fisher Scientific). rGFP-CAAX, rGFP-FYVE, rGFP-Giantin, rGFP-PTP1b, rGFP-NLS, Rap1Gap1a-*R*luc2 plasmid DNA and β-arrestin2-*R*luc2 in the pcDNA3.1(+) backbone were synthesized by GenScript. Venus-KRas, Nluc-mGsi and Venus-mGsi were kind gifts from Nevin Lambert (Augusta University). β-arrestin2-Venus, GRK2-Venus and GRK2-*R*luc2 were kind gifts from Gunnar Schulte (Karolinska Institutet). Plasmid DNA encoding an α subunit of Gi1 was from cDNA.org. Salmon sperm DNA (ss DNA) was from Thermo Fisher Scientific. The constructs were validated by Sanger sequencing (Eurofins GATC).

### Ligand

The endogenous GPR183 agonist 7α,25-dihydroxycholesterol (#SML0541; referred also as the GPR183 agonist throughout the study) was from Sigma. The ligand was dissolved in DMSO at 5 mM, aliquoted, and stored at -20°C. Each aliquot was used a maximum of three times.

### Enhanced bystander BRET (ebBRET; Rluc2-rGFP) setup

HEK293A cells were transiently transfected in suspension using polyethylenimine (PEI, Polysciences). To measure heterotrimeric Gi protein activation/dissociation at the cell membrane interface, a total of ca. 4 × 10^5^ cells were transfected in 1 mL with 200 ng of HiBiT-GPR183 plasmid DNA constructs, 300 ng of rGFP-CAAX plasmid DNA, 100 ng of an α subunit of Gi1 plasmid DNA, 50 ng of Rap1Gap-*R*luc2 plasmid DNA and 350 ng of ssDNA. To measure GRK2 recruitment to the cell membrane-expressed GPR183 in an ebBRET paradigm, a total of ca. 4 × 10^5^ cells in 1 mL were transfected with 200 ng of HiBiT-GPR183 plasmid DNA, 50 ng of GRK2-*R*luc2 plasmid DNA, 300 ng of rGFP-CAAX plasmid DNA and 450 ng of ssDNA. To measure β-arrestin2 recruitment to the cell membrane-expressed GPR183 in an ebBRET paradigm, a total of ca. 4 × 10^5^ cells in 1 mL were transfected with 200 ng of HiBiT-GPR183 plasmid DNA, 50 ng of β-arrestin2-*R*luc2 plasmid DNA, 300 ng of rGFP-CAAX plasmid DNA and 450 ng of ssDNA. Next, transfected cells (4 × 10^4^ cells in 100 μL) were seeded onto white 96-well cell culture plates. 24h later, the cells were washed once with 200 µL of HBSS (HyClone). Next, 80 μL of HBSS was added to the wells, and subsequently, 10 μL of furimazine (1:1000 final concentration, Promega) was added. The plate was incubated for 10 min, which was followed by the addition of 10 μL of the agonist. Next, *R*luc2 emission (donor, 360-440 nm, 100 ms integration time) and rGFP emission (acceptor, 505-575 nm, 100 ms integration time) were measured at 37°C (three measurements for baseline, 15 min with the agonist). The ebBRET ratios were defined as acceptor emission/donor emission. The measurements were performed using a Tecan Spark microplate reader.

### Nluc-Venus BRET setup

HEK293A cells were transiently transfected in suspension using polyethylenimine (PEI, Polysciences). To measure miniGsi recruitment to the receptor, a total of ca. 4 × 10^5^ cells were transfected in 1 mL with 250 ng of Venus-mGsi plasmid DNA construct, 50 ng of HiBiT-GPR183-Nluc plasmid DNA, 700 ng of ssDNA. To measure GRK2 recruitment to the receptor, a total of ca. 4 × 10^5^ cells were transfected in 1 mL with 250 ng of GRK2-Venus plasmid DNA construct, 50 ng of HiBiT-GPR183-Nluc plasmid DNA, 700 ng of ssDNA. To measure β-arrestin2 recruitment to the receptor, a total of ca. 4 × 10^5^ cells were transfected in 1 mL with 250 ng of β-arrestin2-Venus plasmid DNA construct, 50 ng of HiBiT-GPR183-Nluc plasmid DNA, 700 ng of ssDNA. To measure miniGsi recruitment to the receptor at the plasma interface, a total of ca. 4 × 10^5^ cells were transfected in 1 mL with 250 ng of Venus-Kras plasmid DNA construct, 10 ng of Nluc-mGsi plasmid DNA, 100 ng of HiBiT-GPR183 plasmid DNA and 640 ng of ssDNA. To receptor expression at the cell surface, a total of ca. 4 × 10^5^ cells were transfected in 1 mL with 900 ng of Venus-Kras plasmid DNA construct and 100 ng of GPR183-Nluc. Next, transfected cells (4 × 10^4^ cells in 100 μL) were seeded onto white 96-well cell culture plates. 24h later, the cells were washed once with 200 µL of HBSS (HyClone). Next, 80 μL of HBSS was added to the wells, and subsequently, 10 μL of furimazine (1:1000 final concentration, Promega) was added. The plate was incubated for 10 min, which was followed by the addition of 10 μL of the agonist. Next, Nluc emission (donor, 460-500 nm, 50 ms integration time) and Venus emission (acceptor, 520-560 nm, 50 ms integration time) were measured at 37°C (three measurements for baseline, 15 min with the agonist). The BRET ratios were defined as acceptor emission/donor emission. The measurements were performed using a Tecan Spark microplate reader.

### GPR183 conformational sensors experiments

HEK293A cells were transiently transfected in suspension using PEI. To measure ligand-induced conformational change in GPR183 a total of ca. 4 × 10^5^ cells were transfected in 1 mL with 50 ng of HiBiT-GPR183-mNG-Nluc plasmid DNA constructs and 950 ng of ssDNA or 10 ng of HiBiT-GPR183-Halo-Nluc plasmid DNA and 990 ng of ssDNA. Next, transfected cells (4 × 10^4^ cells in 100 μL) were seeded onto white or black 96-well cell culture plates. 24 h later, the cells were washed once with 200 µl of HBSS (HyClone). Next, 80 μL of HBSS was added to the wells, and subsequently, 10 μL of 1:1000 final furimazine (Promega) was added for the mNG-Nluc sensors The plate was incubated for 10 min, which was followed by the ligand Next, Nluc emission (donor, 460-500 nm, 50 ms integration time) and mNG emission (acceptor, 505-545 nm, 50 ms integration time) were measured at 37°C (three measurements for baseline, 15 min with agonis). The BRET ratio were defined as acceptor emission/donor emission. The measurements were performed using a Tecan Spark microplate reader.

### Luminescence-based measurements of cell surface expression of N-terminally HiBiT-tagged receptors

HEK293A cells at a density of 4 x 10^5^ cells/mL were transfected in suspension. 3 x 10^4^ RAW264.7 cells per well of a 96-well plate were transfected 24 h following seeding. HEK293A cells were transfected using PEI with 200 ng of the indicated receptor plasmid DNA (HiBiT-GPR183 wild type (WT) or A338V) and 800 ng ssDNA. HEK293A cells in a volume of 100 μL were seeded onto a white 96-well plate with a flat bottom RAW264.7 cells were transfected using PEI with 20 ng of the indicated receptor plasmid DNA (HiBiT-GPR183 wild type (WT) or A338V) per well. 24 h post-transfection, the cells were washed once in HBSS and incubated with 100 μL of furimazine/LgBiT mix (1:100 final concentration for furimazine and 1:200 final concentration LgBiT, with Promega) for 10 min. Subsequently Nluc/NanoBiT luminescence (460-500 nm, 50 ms integration time) was measured using a Tecan Spark microplate reader.

### Immunoblotting

HEK293A cells were transfected at a density of 4 × 10^5^ cells, with a total volume of 1 mL used per transfection reaction. The DNA mix for each transfection included pcDNA, HiBiT-GPR183 WT or HiBiT-GPR183 A338V. Specifically, the plasmid DNA mixes were composed of 200 ng of receptor plasmid DNA and 800 ng of pcDNA, or just pcDNA for the control. The cells were incubated at 37°C for 24 hours. Whole-cell lysates were prepared by combining 50 µl of 2x Laemmli buffer containing 10% dithiothreitol with 50 µl of PBS. This mixture was subjected to sonication at maximum power in three sequences of 3-second bursts, each separated by 3-second pauses. The lysates were subsequently analyzed using 4–20% Mini-PROTEAN TGX precast polyacrylamide gels (Bio-Rad) along with the PageRuler Plus Prestained Protein Ladder (Thermo Fisher Scientific). The membranes were blocked with 5% non-fat dried milk in a Tris/NaCl/Tween20 buffer (TBS-T) and then incubated overnight at 4°C with primary antibodies diluted in the blocking buffer. The primary antibodies used were rabbit monoclonal anti-GAPDH (14C10; 1:2500 dilution; Cell Signaling Technology #2118) and mouse monoclonal anti-HiBiT (1.0 μg/ml; Promega clone 30E5; IgG2c with kappa light chain). Following several washes with TBS-T, the membranes were incubated with horseradish peroxidase-conjugated secondary antibodies (goat anti-mouse, 1:3000, Thermo Fisher Scientific #31430 and goat anti-rabbit, 1:3000, Thermo Fisher Scientific #31460) diluted in blocking buffer for 1 hour at room temperature. After additional washes, the membranes were exposed to Clarity Western ECL Blotting Substrate (Bio-Rad) for 2 minutes. Protein bands were then visualized using the ChemiDoc system (Bio-Rad) and analysed in ImageLab.

### Statistical analysis

All data presented in the main figures are based on at least three independent experiments (biological replicates), with each individual experiment typically performed at least in duplicates (technical replicates) for each condition, unless otherwise specified in a figure legend. One biological replicate refers to wells containing cells seeded from the same individual cell culture flasks and measured on the same day. Different biological replicates were transfected using separate transfection mixtures and measured on different days. Technical replicates are defined as individual wells with cells from the same biological replicate. Data presented in the Supporting Information, which indeed play a supporting role, may come from fewer than three biological experiments, and furthermore can be presented in the form of a representative example, therefore without a statistical analysis. Samples were not randomized or blinded during the experiments. Statistical and graphical analyses were performed using Graph Pad Prism software (version 10.4.0). Concentration response data were fitted to either a three-parameter or a four-parameter non-linear model selected with the extra sum-of-squares F-test. Two datasets were analysed for statistical differences with paired *t*-test. Three or more datasets were analysed by one-way ANOVA with multiple comparison Dunnett’s post-hoc analysis. Significance levels are given and displayed in the figures as: *P < 0.05; **P < 0.01; ***P < 0.001; ****P < 0.0001. Differences between datasets which did not reach statistical significance are marked with “ns”. Data points throughout the manuscript are indicated as the mean ± standard error of the mean (s.e.m.) unless otherwise stated.

## Results

### The A338V mutation preserves agonist potency but reduces mGsi, GRK2 and β-arrestin2 and engagement in direct BRET assays

To investigate whether the A338V substitution (**Figure 1A)** alters receptor-mediated proximal signaling events, we first performed direct intermolecular BRET assays monitoring recruitment of mGsi (GPR183 is a Gi-coupled receptor), GRK2 and β-arrestin2 to an overexpressed HiBiT-GPR183. WT or A338V GPR183 fused C-terminally to Nluc (GPR183-Nluc) were co-expressed in HEK293A cells with Venus-tagged protein partners: Venus-mGsi, GRK2-Venus or β-arrestin2-Venus. Agonist, 7α,25-dihydroxycholesterol, stimulation induced robust increases in BRET signal in all configurations, confirming functional receptor activation (**Figure 1B–G**). In each case, the maximal agonist-induced BRET response was consistently higher for WT compared to the A338V variant (**Figure 1B–G**). This reduction in BRET suggested decreased efficiency of transducer and mGsi probe engagement in the direct configuration.

**Figure 1.**
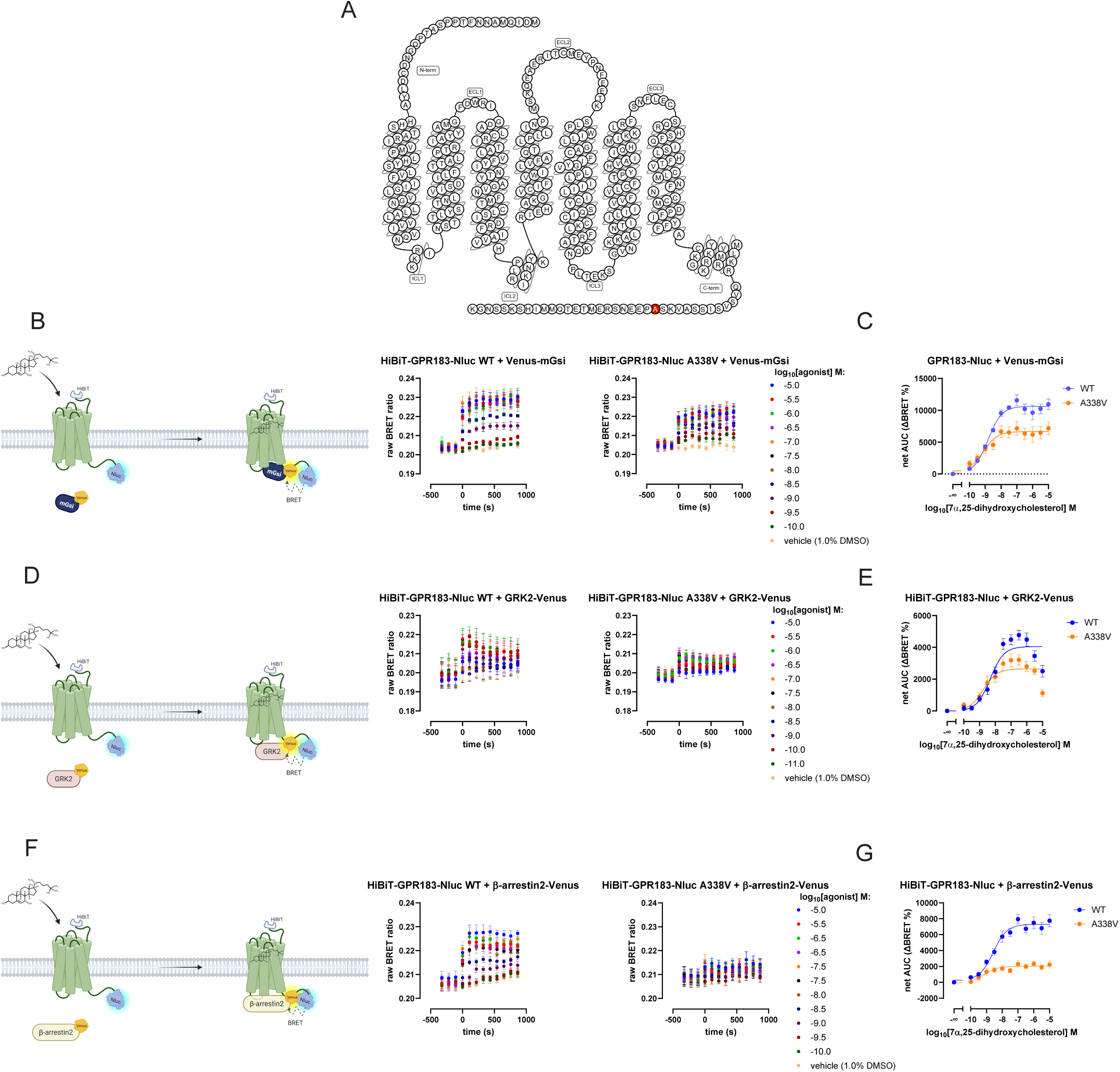
Direct BRET paradigm reveals differences in coupling profiles between GPR183 WT and A338V. **A.** Schematic representation of the human GPR183 with alanine at a position 338 (A338V) shown in red. **B. Left:** Schematic representation of direct BRET-based Venus-mGsi recruitment assay; **Right**: BRET kinetic traces, agonist was added at time = 0. **C.** net AUC representation of the data from the **B right panels. D. Left:** Schematic representation of direct BRET-based GRK2-Venus recruitment assay; **Right**: BRET kinetic traces, agonist was added at time = 0. **E.** net AUC representation of the data from the **D right panels. F. Left:** Schematic representation of direct BRET-based β-arrestin2-Venus recruitment assay; **Right**: BRET kinetic traces, agonist was added at time = 0. **G.** net AUC representation of the data from the **F right panels**. Kinetic traces are represented as mean ± S.E.M. of n≥3 independent experiments, which were performed at least in technical duplicates each; net AUC data are presented with S.D.

Concentration–response curves were fitted to derive *EC*₅₀ values, which did not differ significantly between WT and A338V receptors. Thus, agonist potency sensitivity remained intact in the mutant receptor and the observed differences therefore reflected altered agonist efficacy and coupling efficiency rather than impaired ligand recognition.

Because the A338V substitution resides within the C-terminal tail, 23 amino acids from the distal end - the same region to which Nluc is fused in these constructs - direct BRET measurements may be sensitive to changes in donor–acceptor proximity or dipole orientation. Reduced BRET efficiency in the mutant receptor could theoretically arise from altered spatial positioning of the Nluc donor due to local C-tail conformational changes. We therefore sought to validate these findings using assay configurations that should not in principle depend on C-terminal donor geometry.

### Membrane-recruitment bystander BRET assays confirm preserved Gi activation but impaired GRK2 and β-arrestin2 recruitment

To validate the direct BRET findings, membrane-recruitment assays were performed using HiBiT-tagged GPR183 WT or A338V, rGFP-CAAX (cell membrane marker), and *R*luc2-tagged transducers or effector proteins. This configuration measures BRET between the plasma membrane-expressed protein tagged with rGFP and transducer/effect protein tagged with a BRET donor *R*luc2. To this end, in this paradigm protein interactions between a receptor and its protein binding partner are inferred from and are independent of direct receptor–transducer dipole alignment as in the direct setup.

In the Rap1Gap-*R*luc2-based Gi activation/translocation assay (**Figure 2A**), the A338V variant exhibited higher agonist-induced BRET responses compared to WT (**Figure 2B-C**). However, analysis of basal signaling revealed that A338V displays slightly reduced constitutive Gi activity (**Figure 2D)**. Because agonist responses were analysed in relation to basal levels, the lower baseline resulted in an apparently greater area under the curve (net AUC) for the mutant receptor (**Figure 2C**). When raw agonist-stimulated BRET values were compared directly, WT and A338V responses were largely comparable (**Figure 2B**), indicating preserved total cell Gi activation capacity. Consistent results were obtained in an independent Gi configuration using Nluc-miniGsi and Venus-KRas (**Figure 2E**) with the same HiBiT-tagged receptor constructs, where agonist was equally potent and only slightly more efficacious on the mutant variant (**Figure 2F-G**).

**Figure 2.**
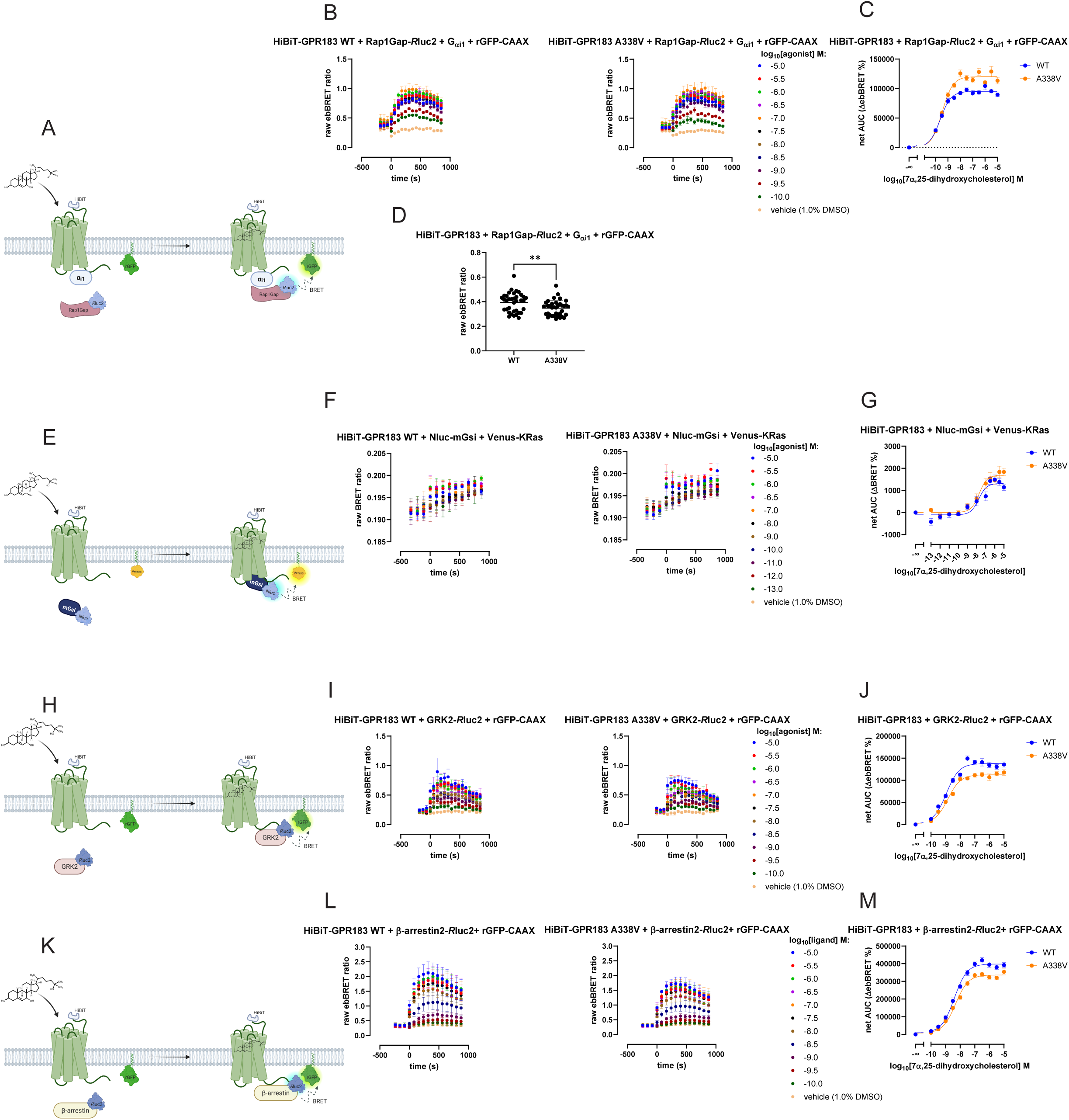
Bystander BRET paradigm demonstrates that GPR183 A338V recruits GRK2 and β-arrestin2 less efficiently than WT. **A.** Schematic representation of ebBRET-based Gi activation assay; **B**. BRET kinetic traces, agonist was added at time = 0. **C.** net AUC representation of the data from **B** panel**. D.** Constitutive activity of HiBiT-GPR183 WT and A338V represented as raw ebBRET values in the two experimental setups prior to ligand stimulation. Each data point represent an individual well from experiments presented in panel A. **E.** Schematic representation of bystander BRET-based Nluc-mGsi recruitment assay. **F.** BRET kinetic traces, agonist was added at time = 0. **G.** net AUC representation of the data from **F** panel**. H.** Schematic representation of ebBRET-based β-arrestin2-*R*luc2 recruitment assay. **I** ebBRET kinetic traces, agonist was added at time = 0. **J.** net AUC representation of the data from **I** panel**. K.** Schematic representation of ebBRET-based GRK2-*R*luc2 recruitment assay. **L.** ebBRET kinetic traces, agonist was added at time = 0. **M.** net AUC representation of the data from **L** panel. Kinetic traces are represented as mean ± S.E.M. of n≥3 independent experiments, which were performed at least in technical duplicates each; net AUC data are presented with S.D.

In contrast, BRET arising from agonist-induced recruitment of GRK2-*R*luc2 and β-arrestin2-*R*luc2 (**Figure 2H and Figure 2K**) was significantly higher for WT compared to A338V (**Figure 2I-M**). Concentration–response analyses revealed similar agonist *EC*₅₀ values between receptor, confirming that ligand potency remains unchanged. These findings suggest that A338V selectively impairs agonist efficacy in the ebBRET assays measuring recruitment of GRK2 and β-arrestin2 while mostly preserving G protein activation.

To determine whether differences in receptor abundance contribute to this phenotype, total and cell surface expression levels of overexpressed WT and A338V GPR183 were assessed in HEK293A cells using the same HiBiT-tagged constructs. Immunoblot analysis revealed stronger total expression of WT compared to A338V (**Figure S1A**), indicating higher overall abundance of the WT receptor. In contrast, cell surface luminescence measurements demonstrated that the A338V variant exhibits increased plasma membrane expression relative to WT (**Figure S1B**). These findings indicate that the reduced GRK2 and β-arrestin2 recruitment observed for A338V cannot be attributed to decreased receptor availability at the cell surface. Furthermore, despite higher membrane expression, the A338V variant displays slightly reduced basal Gi activity, indicating that the mutation affects constitutive signaling independently of receptor abundance. However, absolute agonist-induced Gi activation was comparable between variants. Together, the data support a selective functional effect of the A338V substitution on receptor signaling properties rather than altered receptor expression levels.

### Intramolecular conformational biosensor reveals global activation differences

To determine whether the A338V substitution induces detectable changes in receptor core conformation, we employed an intramolecular conformational biosensor (GPR183-mNG-Nluc, **Figure 3A**). Analysis of basal conformational suggests that for that similar expression levels, basal BRET values were usually higher for the A338V variant, i.e. separate clusters of data points for WT and for A338V can be distinguished in the **Figure 3B**. These data indicate that the mutation does produce measurable alterations in global receptor conformation prior to an agonist activation. However, agonist-induced conformational changes did not differ significantly between WT and A338V receptors (**Figure 3 C-D**). Still, what these data suggest fit well with the results obtained for the ebBRET-based analysis of Gi activation indicating that, prior to the agonist addition, the A338V mutant remains in a more inactive conformation (higher BRET values with the conformational sensor) that allows less efficient activation of Gi (lower ebBRET values in the Gi setup). Nevertheless, it is imperative to consider potential effects of the C-tail mutant on the positioning of the Nluc tag that could play a role in efficiency of energy transfer between Nluc and mNG.

**Figure 3.**
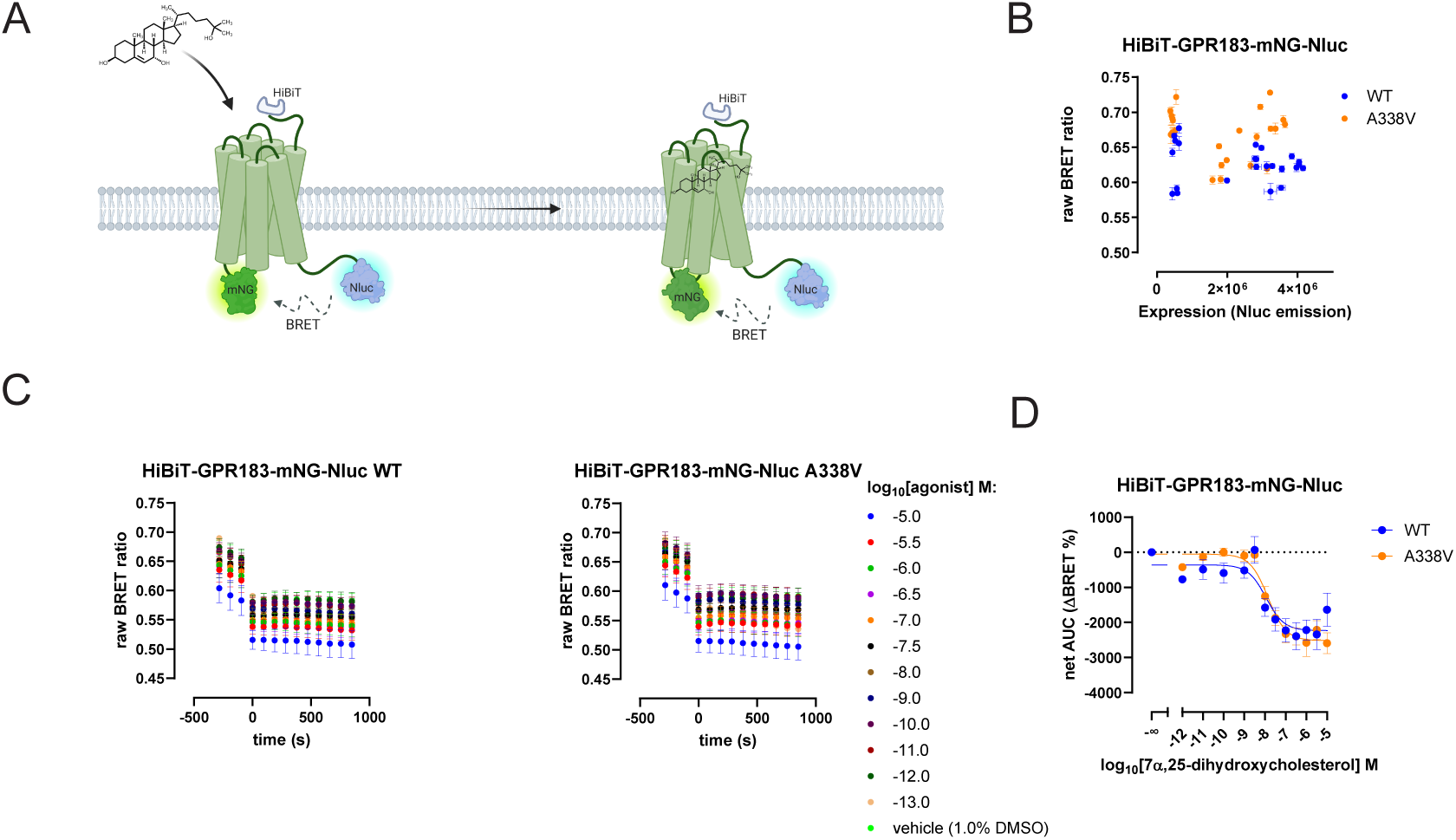
Analysis of conformational dynamics reveals potential differences in basal conformational state of the two variants but no changes in ligand-induced conformation. **A.** Schematic representation of the GPR183 conformational sensor. **B.** raw BRET values obtained for the GPR183-mNG-Nluc WT and A338V for different expression levels. Each data point indicate one well from three separate plates. **C**: BRET kinetic traces in the conformational sensor experiment, agonist was added at time = 0. **D.** net AUC representation of the data from the **C** panels.

### Nluc dark experiments probe C-terminal structural effects

To further investigate the basis of the reduced transducers recruitment observed for A338V, we employed a enzyme-deficient Nluc R164Q construct (Nluc* “Nluc dark”; R164Q denotes mutation in the Nluc) [37, 38] and used it fused with receptors’ C-tail in the already introduced assays (**Figure 4A**). In this configuration, presence of the tag on the receptor does not compromise the measurements as luminescence originates from *R*luc2-tagged Rap1Gap, β-arrestin2 and GRK, or Nluc-tagged miniGsi and not from the receptor, allowing recruitment to be assessed independently of receptor-derived energy transfer while preserving the structural context of the C-terminal Nluc fusion.

**Figure 4.**
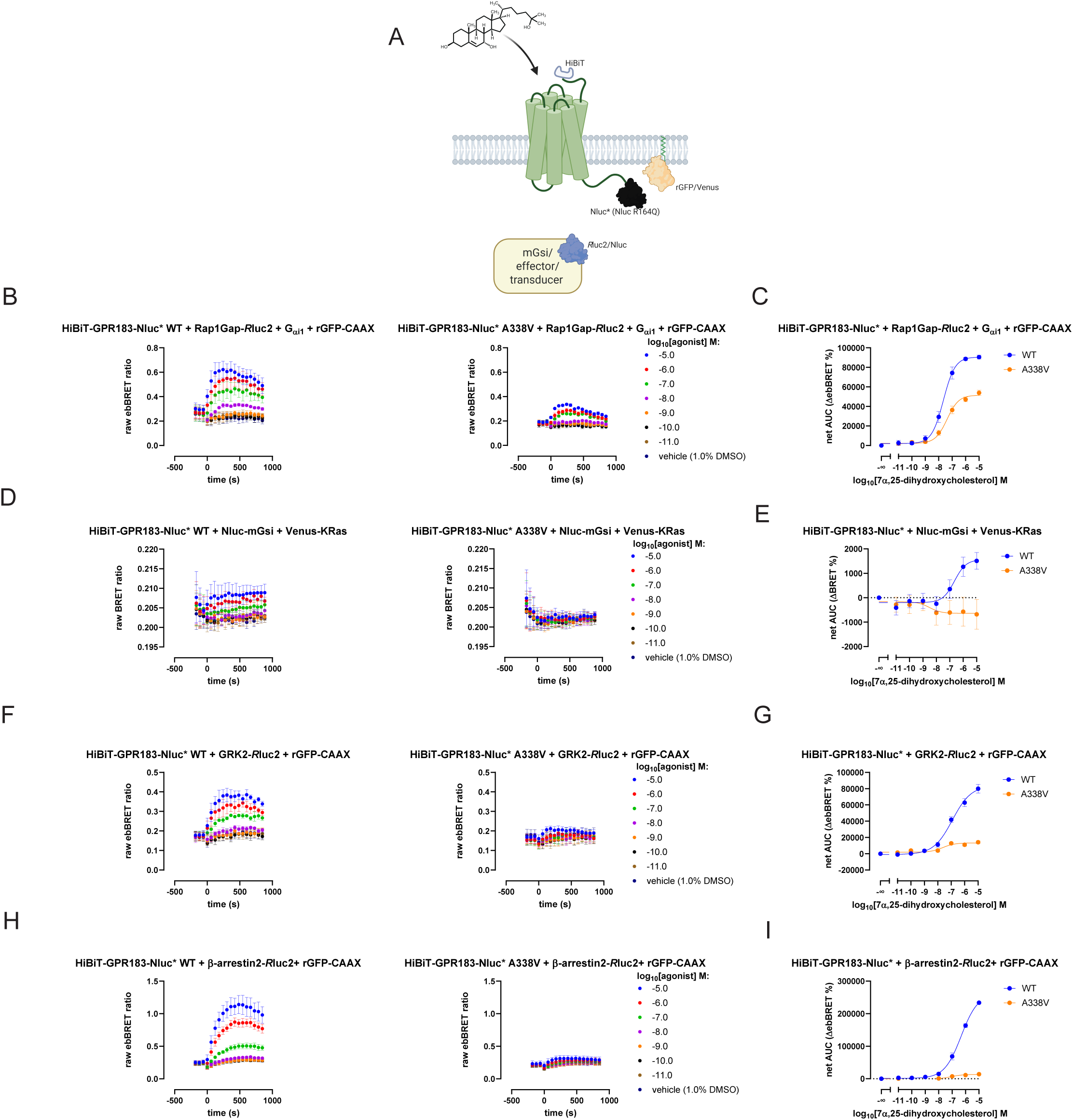
Bystander BRET experiments with C-tail tag receptors indicate that A338V introduces substantial changes in the C-tail conformation. **A. Left:** Schematic representation of different experimental paradigms used with an overexpressed HiBiT-GPR183-Nluc*; **Right**: **B.** ebBRET kinetic traces in an ebBRET-based Gi activation experiment, agonist was added at time = 0. **C.** net AUC representation of the data from the **B. D. Left:** BRET kinetic traces in a bystander BRET-based Nluc-mGsi recruitment, agonist was added at time = 0. **E.** net AUC representation of the data from the **D** panel. **F.** ebBRET kinetic traces in an ebBRET-based GRK2-*R*luc2 recruitment experiment, agonist was added at time = 0. **G.** net AUC representation of the data from the **F panel. H.** ebBRET kinetic traces in an ebBRET-based β-arrestin2-*R*luc2 recruitment experiment, agonist was added at time = 0. **I.** net AUC representation of the data from the **F panel.** ebBRET Kinetic traces are represented as mean ± S.E.M. of n≥3 independent experiments, which were performed at least in technical duplicates each; net AUC data are presented with S.D.

Agonist-induced recruitment of binding partners and Rap1Gap, as a probe for activated Gi, to the membrane bound GPR183 remained significantly reduced for the A338V variant compared to WT (**Figure 4 B-E**), consistent with the membrane recruitment data and, except for the Nluc-mGsi setup, in fact very closely mirroring results obtained in the Nluc-Venus setup. Interestingly, in the Nluc-mGsi setup, the presence of the Nluc on the GPR183 A338V C-tail seems to completely abolish any ligand-stimulated recruitment of this conformational probe, and in fact appears to dissociate it from its initial binding at the overexpressed receptor. Again, since the Nluc dark moiety remains present but does not serve as the luminescent donor, this setup allows discrimination between altered BRET efficiency and genuine changes in recruitment.

The persistence of reduced Gi activation, as well as diminished miniGsi, β-arrestin2, GRK2 engagement with the ligand-stimulated A338V in the Nluc dark configuration indicates that the results are not solely attributable to donor–acceptor geometry but are likely to arise from steric interference caused by 17 kDa Nluc tag. It needs to be noted here, that the A338V mutant of the GPR183 fused to Nluc R164Q had significantly lower cell surface expression levels, which could contribute to reduced observed ligand-activated receptor-mediated phenomena, but should not abolish them as the expression levels of the A388V construct were still substantial (**Figure S2A**). Along these lines, an orthogonal measurement of cell surface expression employing GPR183-Nluc WT or A338V together with Venus-Kras also resulted in statistically significant differences in cell surface expression between the WT and A338V (**Figure S2B)**. However, here the differences in BRET can also arise from different dipole-dipole orientations and/or distance between the BRET pair supporting different tag orientation as a result of the C-tail mutation. Overall, these findings support the interpretation that the A338V substitution alters the conformational properties or spatial accessibility of the C-terminal tail itself, thereby limiting productive intracellular transducer coupling to an overexpressed GPR183 A338V.

### MD simulations

MD simulations reveal that the A338V mutation substantially reorganises the conformational landscape of GPR183, primarily through structural rearrangements in the C-terminal tail (C-tail). From the trajectories obtained for each system (the active and inactive structures were used as starting pose for the simulations of GPR183 without the Nluc tag), five representative structural clusters were selected and are shown in the Supporting Information (**Figure S3**), together with disconnectivity graphs derived from these trajectories (**Figure S4-S9**) illustrating the topology of the underlying energy landscapes. Comparison of the WT and mutant systems indicates that the most pronounced differences arise from conformational changes within the C-tail, suggesting that this region plays a central role in mediating the structural effects of the mutation.

The C-terminal tail exhibits strong activation-dependent structural changes that are further amplified by the A338V mutation after comparing simulations starting from active and inactive state. DSSP-derived secondary-structure probabilities show that the C-tail of GPR183 is highly dynamic and undergoes pronounced conformational rearrangements upon receptor activation (**Figure 5A**). In the inactive state, residues 325–335 display relatively low disorder probabilities, suggesting the presence of a transiently structured segment in the proximal portion of the tail. By contrast, the distal C-terminus (residues ∼340–360) becomes strongly disordered in the active receptor, consistent with activation-dependent release of the tail into a flexible conformation. While the A338V mutation has little effect on the disorder profile of the inactive receptor, it increases disorder probabilities in the downstream region of the tail in the active state. This behaviour indicates that the mutation enhances activation-induced flexibility rather than locally destabilising the structure at the mutation site. Interestingly, comparison with the Nluc-fused constructs further shows that the luciferase tag has only a minor effect on the overall disorder profile of the C-tail.

**Figure 5.**
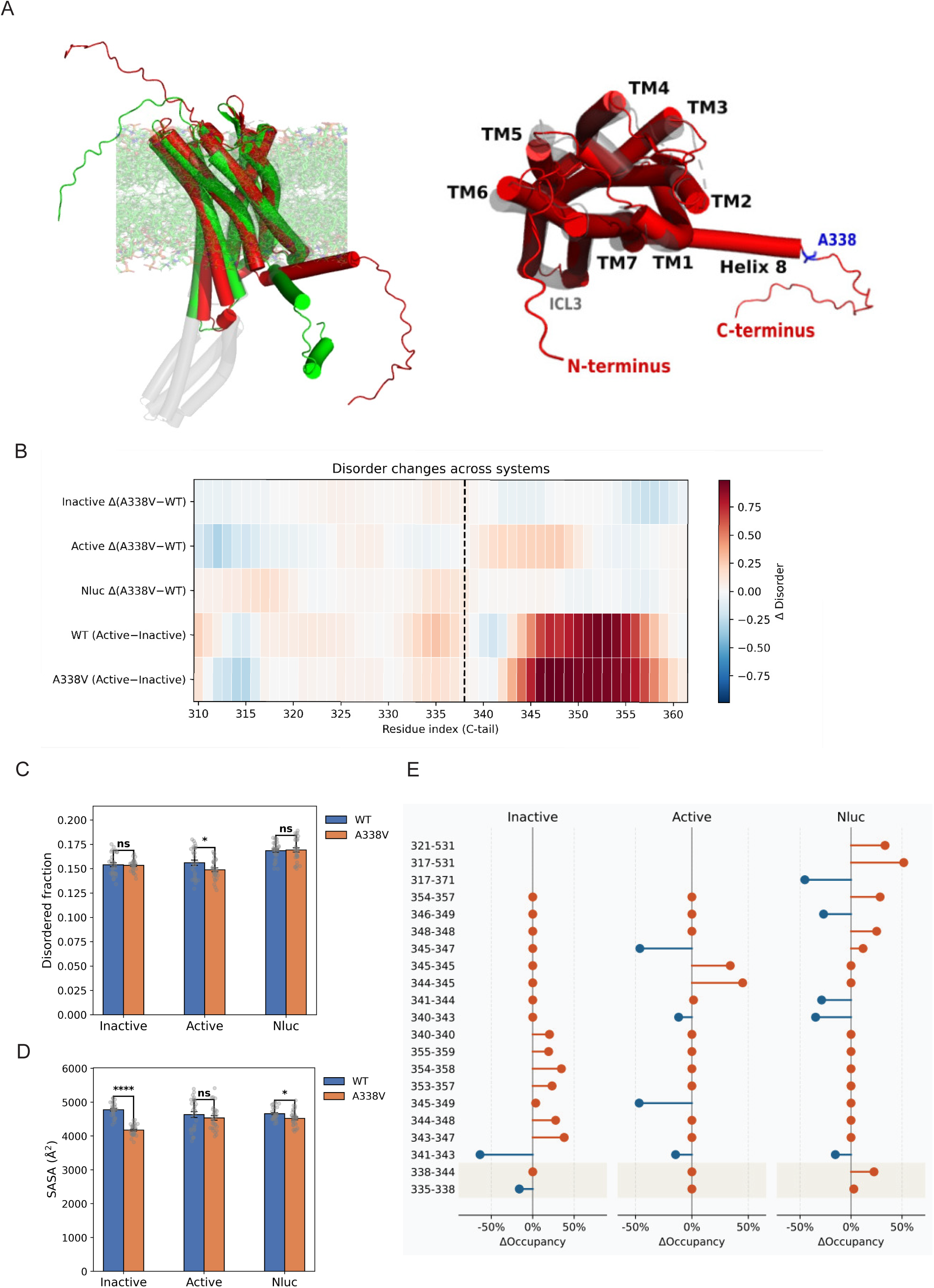
A338V reshapes the C-terminal hydrogen-bonding network, supporting a mutation-driven shift in tail disorder and conformational dynamics. Structural and dynamic effects of the A338V mutation on the GPR183 C-terminal tail. **A.** Structural overview of the receptor systems. Left: schematic representation of the experimental cryo-EM structure together with reconstructed inactive (green) and active (red) conformations of GPR183. Right: intracellular view highlighting the transmembrane helices (TM), intracellular loops (ICL), mutated residue 338, and the N- and C-termini. **B.** Residue-resolved disorder probabilities for the C-terminal tail derived from DSSP analysis across the simulated systems with residue 338 shown with dashed line. **C.** Distribution of the fraction of disordered residues for the full receptor across simulations. **D.** Distribution of solvent-accessible surface area (SASA) of the C-terminal tail for all simulated systems. **E** Differences in C-tail hydrogen-bond occupancies upon the A338V mutation across the analysed systems with H bonds.

Despite increasing disorder locally in the C-tail, the A338V mutation redistributes conformational flexibility across the receptor. To assess global structural variability, we quantified the fraction of disordered residues across the entire receptor in three functional states: inactive, active, and Nluc-fused (**Figure 5C**). In the inactive and Nluc-fused systems, the disorder fraction is similar for the WT and the A338V mutant, with no statistically significant differences. In contrast, the active state shows a modest but statistically significant reduction in the overall disorder fraction for the A338V variant relative to WT, despite the increased disorder observed locally in the C-tail (**Figure 5C**). This behaviour suggests that the mutation does not uniformly increase structural disorder but instead redistributes conformational flexibility within the receptor.

Consistent with these observations, the A338V mutation promotes a more compact conformational ensemble of the C-terminal tail. Analysis of the solvent-accessible surface area (SASA) shows a general reduction in C-tail exposure in the mutant (**Figure 5D**), indicating enhanced self-screening of the tail. This effect is most pronounced in the inactive state, suggesting that the mutation stabilises more compact C-terminal conformations that may influence receptor activation. In the active ensemble the effect is less pronounced, whereas the Nluc-fused system exhibits a statistically significant reduction in C-tail SASA consistent with enhanced compaction.

Hydrogen-bond analysis further demonstrates that A338V reorganises the hydrogen-bonding network of the C-terminal region (**Figure 5E**). In both inactive and active ensembles, hydrogen bonds involving residue 338 decrease in occupancy upon mutation. In the inactive state, the interaction between residues 335–338 decreases by ∼9%, indicating weakened local contacts around the mutation site, while the decrease is smaller in the active ensemble. Beyond the immediate mutation site, however, A338V redistributes hydrogen bonding throughout the C-terminal segment. Several interactions involving residues 340–360 increase markedly in the mutant, including S343–T347, H354–S358, and I353–S357, whereas native contacts such as Q341–S343 show reduced occupancy. These changes indicate a reorganisation of the hydrogen-bonding network rather than a simple loss of interactions.

The mutation also introduces new long-range interactions that alter the spatial positioning of the C-tail. In the inactive ensemble, the mutant forms hydrogen bonds linking the C-terminus with distal regions of the receptor, including contacts involving R317. These interactions are absent or substantially weaker in the WT and may stabilise inactive C-terminal conformations that hinder structural rearrangements associated with activation. Hydrogen bonds involving the Nluc reporter further reveal that A338V promotes additional contacts between the C-terminal segment and luciferase residues, most prominently with A531, whose occupancy increases by up to ∼52%. Concurrently, several native C-terminal hydrogen bonds are weakened or lost.

Collectively, these results demonstrate that A338V reshapes the structural ensemble of the GPR183 C-terminal tail, promoting increased activation-dependent flexibility together with enhanced compaction and a reorganised hydrogen-bonding network. Such large-scale changes in C-tail organisation may alter its interactions with downstream signalling or regulatory partners, providing a structural basis for the functional effects of the mutation.

## Discussion

The present study combines a comprehensive set of cellular assays with molecular simulations to elucidate the functional consequences of the leukaemia-associated A338V mutation in GPR183. The wet-lab data show that this substitution produces selective and pathway-specific functional effects rather than a global disruption of receptor activity, whereas the molecular dynamics simulations provide a structural framework for understanding how these effects may arise.

A key finding from the experimental data is that the A338V mutation preserves agonist potency and overall Gi signaling capacity, as evidenced by both direct and membrane-recruitment BRET assays. Although basal Gi activity was modestly reduced, agonist-stimulated Gi activation remained comparable to the WT receptor across multiple assay configurations. This indicates that the mutation does not impair ligand recognition or the ability of the receptor core to adopt an active conformation capable of engaging G proteins. Consistent with these, intramolecular conformational biosensor experiments seemed to detect differences in global receptor conformation, which could be a component of both differences in the TM3 as well as C-tail conformations, between WT and A338V. Ligand-induced experiments applying the intramolecular conformational sensor showed no differences in the ligand-induced conformational changes, supporting the conclusion that the ligand-induced transmembrane activation mechanism remains largely intact.

Next, the other set of wet-lab experiments consistently revealed a marked reduction in GRK2 and β-arrestin2 recruitment for the A338V variant. This effect was observed across both direct BRET and ebBRET assays and persisted in the Nluc-dark R164Q configuration. Together, these orthogonal approaches strongly indicate that the observed reduction in transducer engagement reflects a genuine functional consequence of the mutation rather than a technical artefact. Importantly, this impairment occurred despite increased plasma membrane expression of the mutant receptor (without the C-tail fusion), excluding reduced receptor availability as an explanation and instead pointing to altered receptor–transducer coupling efficiency. Along these lines, GRK2 and β-arrestin2 interact largely with GPCRs C-tail [12, 39] and as such a different, less accessible conformation of this domain in GPR183 A338V may explain reduced engagement of both proteins.

Functionally then, the receptor remains equally competent for Gi activation but is substantially more deficient in the recruitment of GRK2 and β-arrestin2, indicating that the mutation introduces pathway bias.

The molecular dynamics simulations provide a structural framework for interpreting these experimental observations. The A338V substitution induces widespread reorganisation of the C-terminal tail, including increased activation-dependent disorder, reduced solvent exposure indicative of tail compaction, and a redistribution of hydrogen-bonding interactions. Notably, the mutation promotes the formation of alternative intra-tail and long-range contacts, including interactions linking the C-terminus to other intracellular regions. These changes suggest that the mutation reshapes the conformational ensemble and spatial positioning of the C-tail, rather than causing a simple local perturbation.

The results further indicate that A338V modifies the activation-dependent dynamics of the C-terminal region. In the WT receptor, activation is associated with increased disorder of the distal C-terminal tail, consistent with release of this region into a more flexible conformational state. This behaviour is in line with previous studies showing that intrinsically disordered regions often become more prominent during GPCR activation, facilitating conformational adaptability and signalling interactions [18]. The mutation amplifies this behaviour by increasing disorder in the downstream portion of the tail while simultaneously promoting compaction of the C-terminal ensemble. Such a redistribution of structural flexibility suggests that the mutation alters the dynamic coupling between the receptor core and the regulatory C-terminal region.

Importantly, the structural changes observed in silico align closely with the functional phenotype measured in cells. The C-terminal tail of GPCRs is a critical platform for GRK phosphorylation and β-arrestin binding. The increased compaction and altered hydrogen-bonding network observed for A338V are likely to reduce accessibility or alter the presentation of interaction motifs required for efficient recruitment of regulatory transducers. At the same time, the preservation of global receptor activation and Gi coupling is consistent with the absence of major structural perturbations within the transmembrane core, as supported by both simulations and conformational biosensor data.

The Nluc-based experiments further suggest that the mutation may influence the spatial relationship between the C-tail and fused reporter domains, reflecting altered tail positioning. However, the persistence of reduced recruitment in configurations independent of receptor-derived luminescence argues that steric or conformational constraints imposed by the mutated tail itself are the primary driver of the observed phenotype, rather than tag-related artefacts.

This selective signalling profile may have implications for disease contexts such as acute myeloid leukaemia, where altered receptor regulation rather than complete loss or gain of function could influence cellular behaviour. More broadly, the results highlight the importance of intrinsically disordered regions in GPCR signalling and demonstrate how subtle mutations within these regions can produce biased functional outcomes through modulation of conformational ensembles rather than discrete structural changes. The simulations also suggest that the mutation can influence the spatial positioning of the C-terminal region relative to the reporter domain in the Nluc constructs. The formation of additional hydrogen bonds between the C-tail and luciferase residues in the mutant indicates increased proximity and structural coupling between these regions. These structural changes provide a plausible explanation for the altered behaviour observed in the BRET-based experiments, as changes in the conformational ensemble of the C-tail would be expected to influence the relative orientation and distance between the receptor and the reporter domain.

Taken together, the results indicate that the A338V mutation primarily causes reshaping the structural ensemble of the GPR183 C-terminal tail, rather than inducing a simple local destabilisation at the mutation site. Because the C-terminal region of GPCRs often mediates interactions with signalling and regulatory partners, such changes in conformational organisation may influence receptor coupling and downstream signalling. The observed mutation-induced redistribution of flexibility and hydrogen-bonding interactions therefore provides a mechanistic framework for understanding how a single residue substitution in the C-tail can propagate structural effects across the receptor and modulate its functional behaviour.

## Study limitations

Several limitations of this study should be noted. First, we focused on the immediate steps in the receptor activation pathway and therefore the impact of the A338V on the GPR183-induced phenotypic changes and any potential role in AML is yet to be established. Second, the structural conclusions are derived from atomistic molecular dynamics simulations and therefore reflect the behaviour of the modelled receptor systems rather than direct experimental structural observations. Although the simulations provide extensive sampling of the conformational landscape, the accessible timescales remain limited relative to the full range of GPCR dynamics. In addition, the systems do not explicitly include signalling partners such as G proteins or arrestins, which may further influence the organisation of the C-terminal tail in the cellular context. However, we do plan to extend analysis to involve these interactions in longer simulations in future work. Finally, while the Nluc fusion construct enables interpretation of the experimental BRET measurements, the reporter domain may impose additional constraints on C-tail dynamics. Despite these limitations, the results consistently indicate that the A338V mutation induces substantial reorganisation of the C-terminal conformational ensemble.

## Supporting Information

### Supplementary figures legends

**Supplementary Figure 1. GPR183 A338V is very efficiently trafficked to the cell membrane despite lower total cell expression than WT. A.** Immunoblotting data showing total cell expression of HiBiT-GPR183 WT or A338V. Membranes were stained with anti-HiBiT and anti-GAPDH antibodies. The data presented here come from a representative experiment from three independent experiments. Predicted molecular weight of HiBiT-GPR183 = 42.5 kDa. **B.** Cell surface expression of HiBiT-GPR183 WT or A338V in live cells. The data are represented as mean ± S.E.M. of n=3 independent experiments; pcDNA-transfected cells were used as a control.

**Supplementary Figure 2. GPR183-Nluc* A338V is less expressed at the cell membrane than the WT. A.** Cell surface expression of HiBiT-GPR183-Nluc* WT or A338V in live cells. The data are represented as mean ± S.E.M. of n=3 independent experiments. **B. Left:** Schematic representation of the experimental setup to assess cell surface expression by measuring BRET between the C-terminally Nluc-tagged receptor and cell membrane-inserted Venus. This setup would also measure differences in Nluc tag positioning in relation to the fixed Venus. **Right:** BRET data from experiments where the HEK293A cells were transfected with fixed amounts of the receptor plasmid DNA construct and Venus-KRas construct. The data are represented as mean ± S.E.M. of n=3 independent experiments; Nluc-DVL2 only was used as a control to measure background BRET in the absence of an energy acceptor.

**Supplementary Figure 3. The cluster populations highlight differences in the conformational ensembles across systems and mutation states.** Five most populated structural clusters extracted from the MD trajectories for each system. Representative structures are shown together with their relative populations within the simulated conformational ensembles.

**Supplementary Figure 4-9. A338V alters the distribution along the C-terminal extension coordinate, indicating a mutation-driven reweighting of compact and extended conformational basins.** Disconnectivity graphs constructed from representative MD trajectories for each simulated system, shown together with the corresponding potential energy traces. The graphs are coloured according to an order parameter defined as the distance between the centre of the vector extending from helix 8 to the end of the C-terminal tail. This metric reflects the degree of C-terminal extension, allowing compact and extended conformations sampled along the trajectories to be distinguished**. Supplementary Figure 4**. Active GPR183 WT, **Supplementary Figure 5**. Active GPR183 A338V mutant, **Supplementary Figure 6**. Inactive GPR183 WT, **Supplementary Figure 7**. Inactive GPR183 A338V mutant, **Supplementary Figure 8**. GPR183-Nluc wild type, **Supplementary Figure 9**. GPR183-Nluc A338V.

## Acknowledgments

P.K. acknowledges support from Gunnar Schulte.

## Data availability

All data sets generated and analysed during the current study are available from the corresponding authors on reasonable request.

## Abbreviations

GPCR: G protein-coupled receptor
MD: molecular dynamics
BRET: bioluminescence resonance energy transfer
ebBRET: enhanced bystander bioluminescence resonance energy transfer

## References

1. Lorente, J.S., et al., GPCR drug discovery: new agents, targets and indications. Nat Rev Drug Discov, 2025.

2. Hauser, A.S., et al., GPCR activation mechanisms across classes and macro/microscales. Nat Struct Mol Biol, 2021. 28(11): p. 879–888.

3. Zhou, Q., et al., Common activation mechanism of class A GPCRs. Elife, 2019. 8.

4. Liu, C., et al., Oxysterols direct B-cell migration through EBI2. Nature, 2011. 475(7357): p. 519-23.

5. Hannedouche, S., et al., Oxysterols direct immune cell migration via EBI2. Nature, 2011. 475(7357): p. 524-7.

6. Kjaer, V.M.S., et al., Migration mediated by the oxysterol receptor GPR183 depends on arrestin coupling but not receptor internalization. Sci Signal, 2023. 16(779): p. eabl4283.

7. Ribeiro, M.L., et al., G protein-coupled receptor 183 mediates the sensitization of Burkitt lymphoma tumors to CD47 immune checkpoint blockade by anti-CD20/PI3Kdeltai dual therapy. Front Immunol, 2023. 14: p. 1130052.

8. Zhuang, H., et al., High Expression of GPR183 Predicts Poor Survival in Cytogenetically Normal Acute Myeloid Leukemia. Biochem Genet, 2025.

9. Ley, T.J., et al., DNA sequencing of a cytogenetically normal acute myeloid leukaemia genome. Nature, 2008. 456(7218): p. 66-72.

10. Ding, L., et al., Clonal evolution in relapsed acute myeloid leukaemia revealed by whole-genome sequencing. Nature, 2012. 481(7382): p. 506-10.

11. Welch, J.S., et al., The origin and evolution of mutations in acute myeloid leukemia. Cell, 2012. 150(2): p. 264–78.

12. Lobbert, A., et al., GPCR kinases phosphorylate GPCR C-terminal peptides in a hierarchical manner. Commun Biol, 2025. 8(1): p. 899.

13. Bertalovitz, A.C., et al., Frizzled-4 C-terminus Distal to KTXXXW Motif is Essential for Normal Dishevelled Recruitment and Norrin-stimulated Activation of Lef/Tcf-dependent Transcriptional Activation. J Mol Signal, 2016. 11: p. 1.

14. Kim, K.M. and M.G. Caron, Complementary roles of the DRY motif and C-terminus tail of GPCRS for G protein coupling and beta-arrestin interaction. Biochem Biophys Res Commun, 2008. 366(1): p. 42–7.

15. Gusach, A., J. Garcia-Nafria, and C.G. Tate, New insights into GPCR coupling and dimerisation from cryo-EM structures. Curr Opin Struct Biol, 2023. 80: p. 102574.

16. Sala, D., P.W. Hildebrand, and J. Meiler, Biasing AlphaFold2 to predict GPCRs and kinases with user-defined functional or structural properties. Front Mol Biosci, 2023. 10: p. 1121962.

17. Hilger, D., The role of structural dynamics in GPCR-mediated signaling. FEBS J, 2021. 288(8): p. 2461–2489.

18. Xu, J., et al., The Role of Intrinsically Disordered Domains in Regulating G Protein-Coupled Receptor Signaling. J Am Chem Soc, 2026. 148(1): p. 1430–1443.

19. Venkatakrishnan, A.J., et al., Structured and disordered facets of the GPCR fold. Curr Opin Struct Biol, 2014. 27: p. 129–37.

20. Chen, H., W. Huang, and X. Li, Structures of oxysterol sensor EBI2/GPR183, a key regulator of the immune response. Structure, 2022. 30(7): p. 1016–1024 e5.

21. Abramson, J., et al., Accurate structure prediction of biomolecular interactions with AlphaFold 3. Nature, 2024. 630(8016): p. 493-500.

22. Fiser, A. and A. Sali, ModLoop: automated modeling of loops in protein structures. Bioinformatics, 2003. 19(18): p. 2500–1.

23. Mirdita, M., et al., ColabFold: making protein folding accessible to all. Nat Methods, 2022. 19(6): p. 679–682.

24. Pieper, U., et al., ModBase, a database of annotated comparative protein structure models and associated resources. Nucleic Acids Res, 2014. 42(Database issue): p. D336-46.

25. Schott-Verdugo, S. and H. Gohlke, PACKMOL-Memgen: A Simple-To-Use, Generalized Workflow for Membrane-Protein-Lipid-Bilayer System Building. J Chem Inf Model, 2019. 59(6): p. 2522–2528.

26. Mark, P. and L. Nilsson, Structure and dynamics of the TIP3P, SPC, and SPC/E water models at 298 K. Journal of Physical Chemistry B, 2001. 105(43): p. 24a-24a.

27. Maier, J.A., et al., ff14SB: Improving the Accuracy of Protein Side Chain and Backbone Parameters from ff99SB. J Chem Theory Comput, 2015. 11(8): p. 3696–713.

28. Dickson, C.J., et al., Lipid14: The Amber Lipid Force Field. J Chem Theory Comput, 2014. 10(2): p. 865–879.

29. Ciura, P., et al., Multilayered Computational Framework for Designing Peptide Inhibitors of HVEM-LIGHT Interaction. J Phys Chem B, 2024. 128(28): p. 6770–6785.

30. Roe, D.R. and T.E. Cheatham, 3rd, PTRAJ and CPPTRAJ: Software for Processing and Analysis of Molecular Dynamics Trajectory Data. J Chem Theory Comput, 2013. 9(7): p. 3084–95.

31. Neuman, V., et al., Visualizing the energy landscape for a molecular dynamics trajectory. J Chem Phys, 2026. 164(4).

32. Becker, O.M. and M. Karplus, The topology of multidimensional potential energy surfaces: Theory and application to peptide structure and kinetics. Journal of Chemical Physics, 1997. 106(4): p. 1495–1517.

33. Wales, D.J., M.A. Miller, and T.R. Walsh, Archetypal energy landscapes. Nature, 1998. 394(6695): p. 758-760.

34. Savitzky, A. and M.J.E. Golay, Smoothing + Differentiation of Data by Simplified Least Squares Procedures. Analytical Chemistry, 1964. 36(8): p. 1627-&.

35. Kozielewicz, P., et al., Quantitative Profiling of WNT-3A Binding to All Human Frizzled Paralogues in HEK293 Cells by NanoBiT/BRET Assessments. ACS Pharmacology & Translational Science, 2021. 4(3): p. 1235–1245.

36. Kozielewicz, P., et al., Structural insight into small molecule action on Frizzleds. Nat Commun, 2020. 11(1): p. 414.

37. Xiong, Y., et al., Engineered Amber-Emitting Nano Luciferase and Its Use for Immunobioluminescence Imaging In Vivo. J Am Chem Soc, 2022. 144(31): p. 14101–14111.

38. Bowin, C.F., et al., WNT stimulation induces dynamic conformational changes in the Frizzled-Dishevelled interaction. Sci Signal, 2023. 16(779): p. eabo4974.

39. Haider, R.S., et al., beta-arrestin1 and 2 exhibit distinct phosphorylation-dependent conformations when coupling to the same GPCR in living cells. Nat Commun, 2022. 13(1): p. 5638.

