## Supplementary figures and images for "Decoding the Structural and Functional Impact of the Leukaemia-Associated A338V Mutation in GPR183"

### Supplementary Figure 1

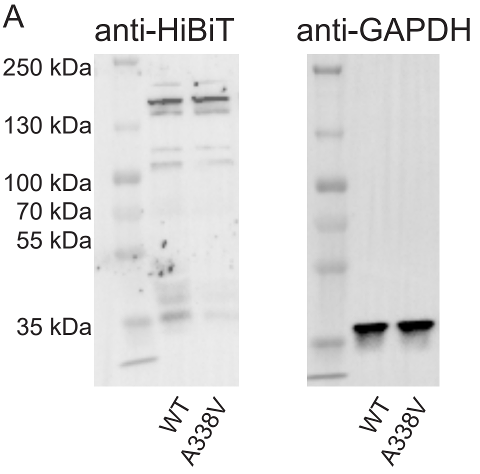

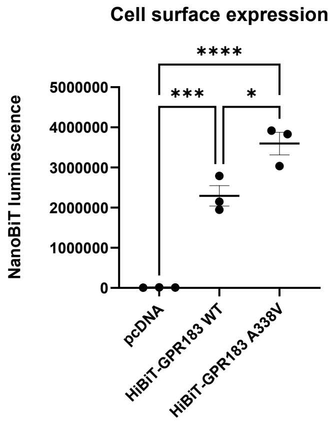

### Supplementary Figure 2

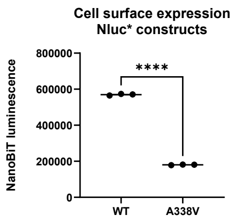

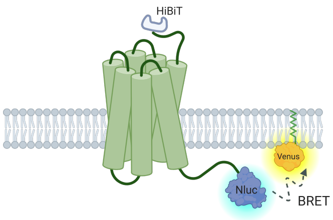

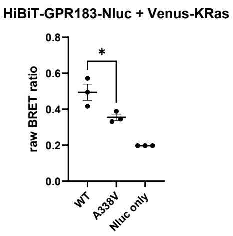

### Supplementary Figure 3

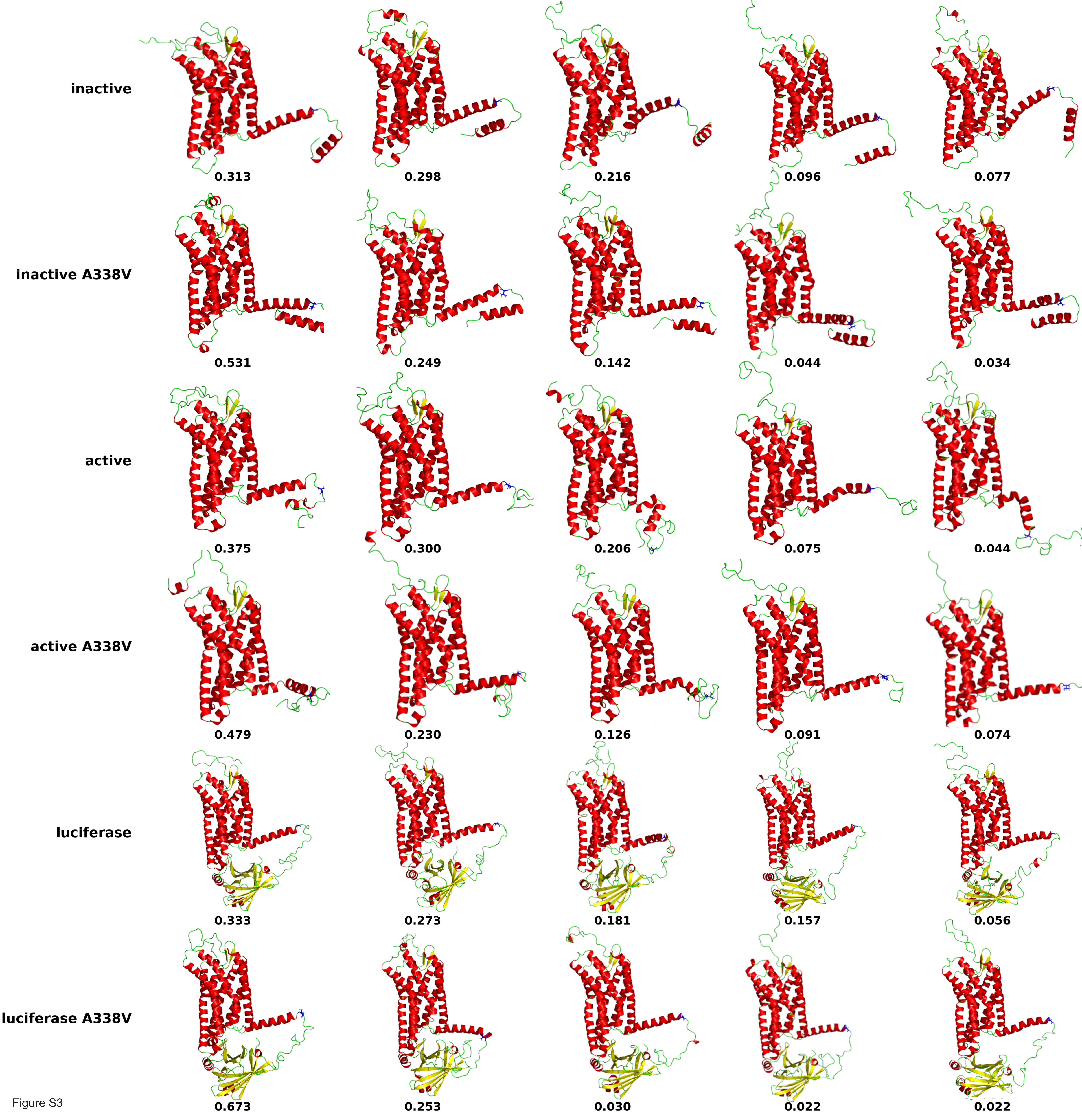

### Supplementary Figure 4

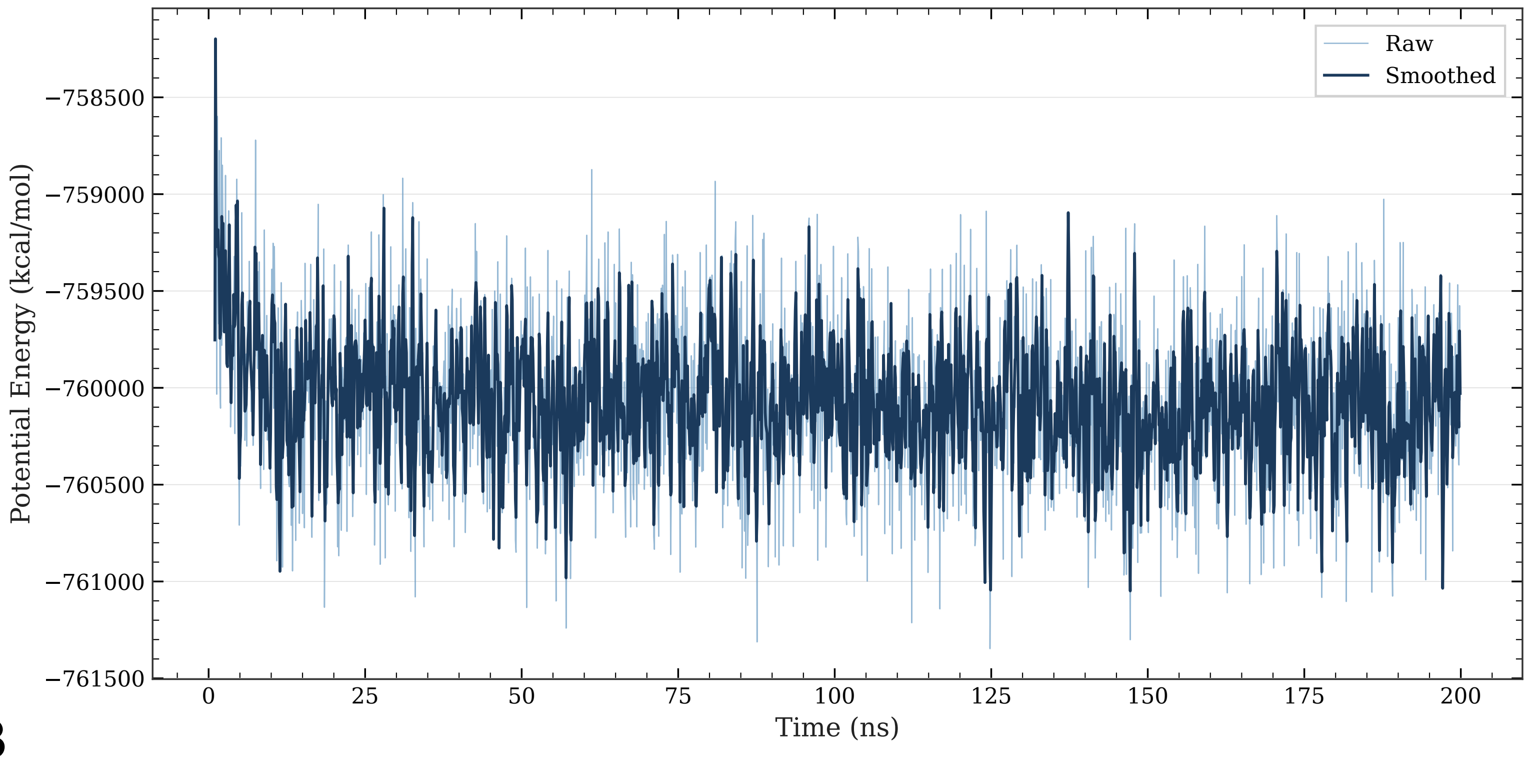

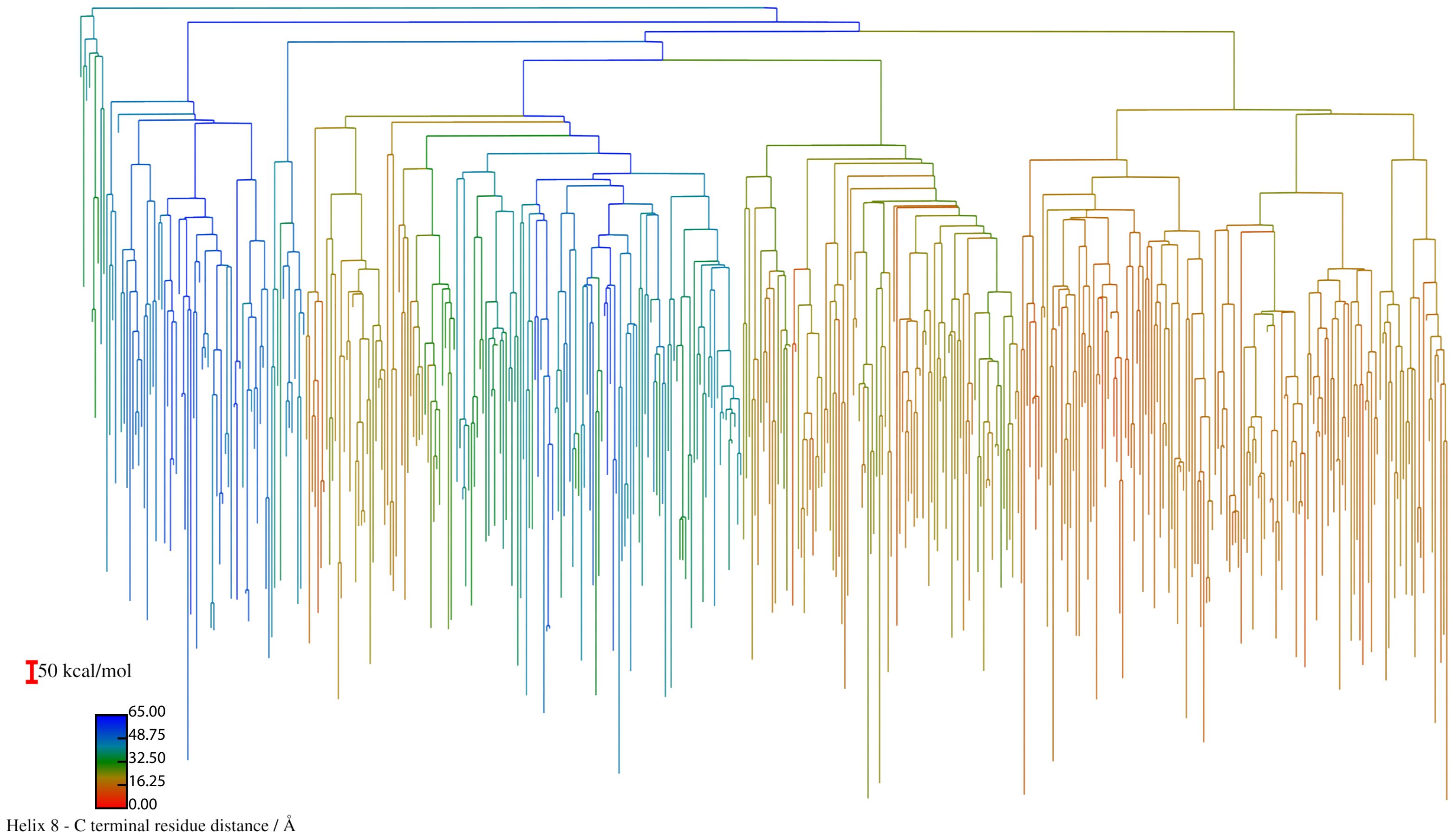

### Supplementary Figure 5

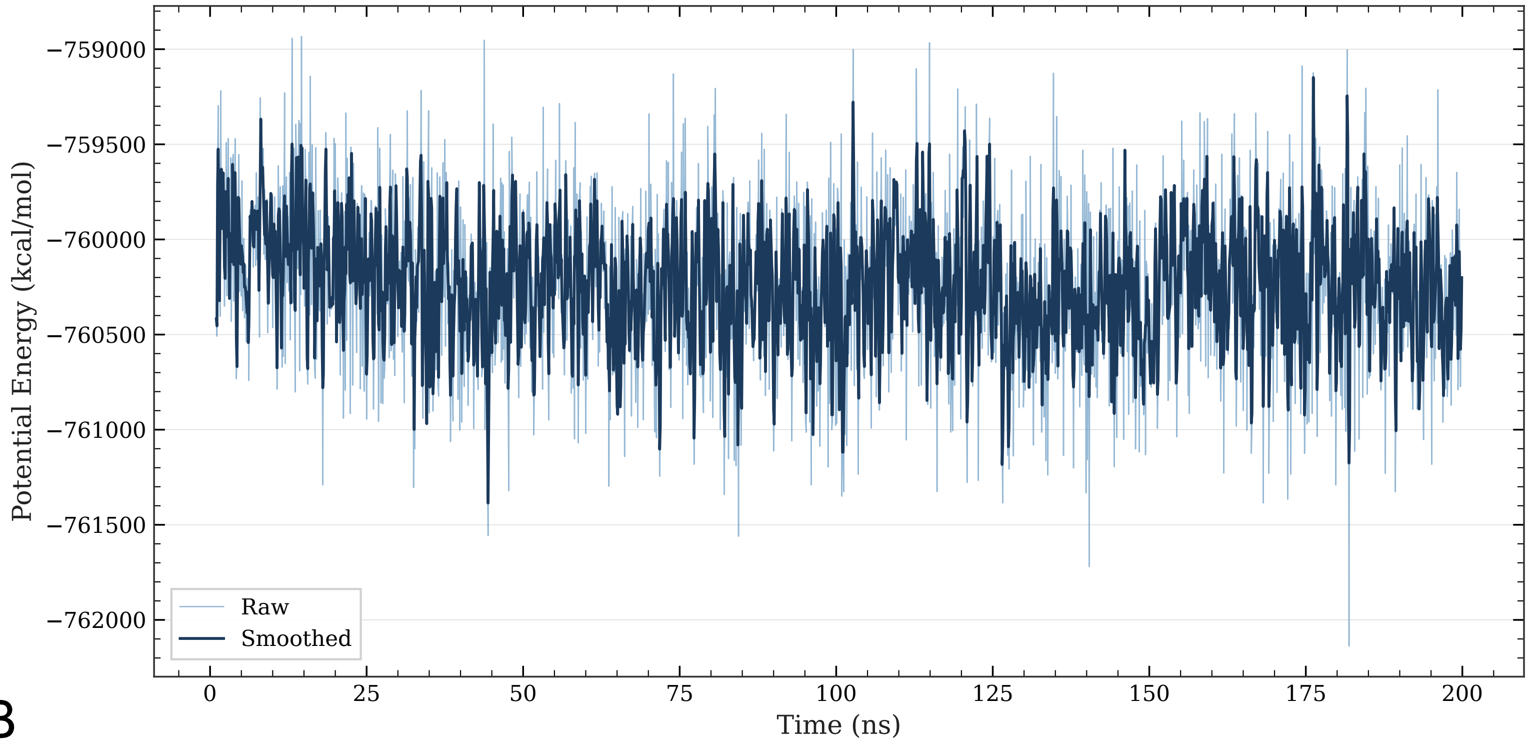

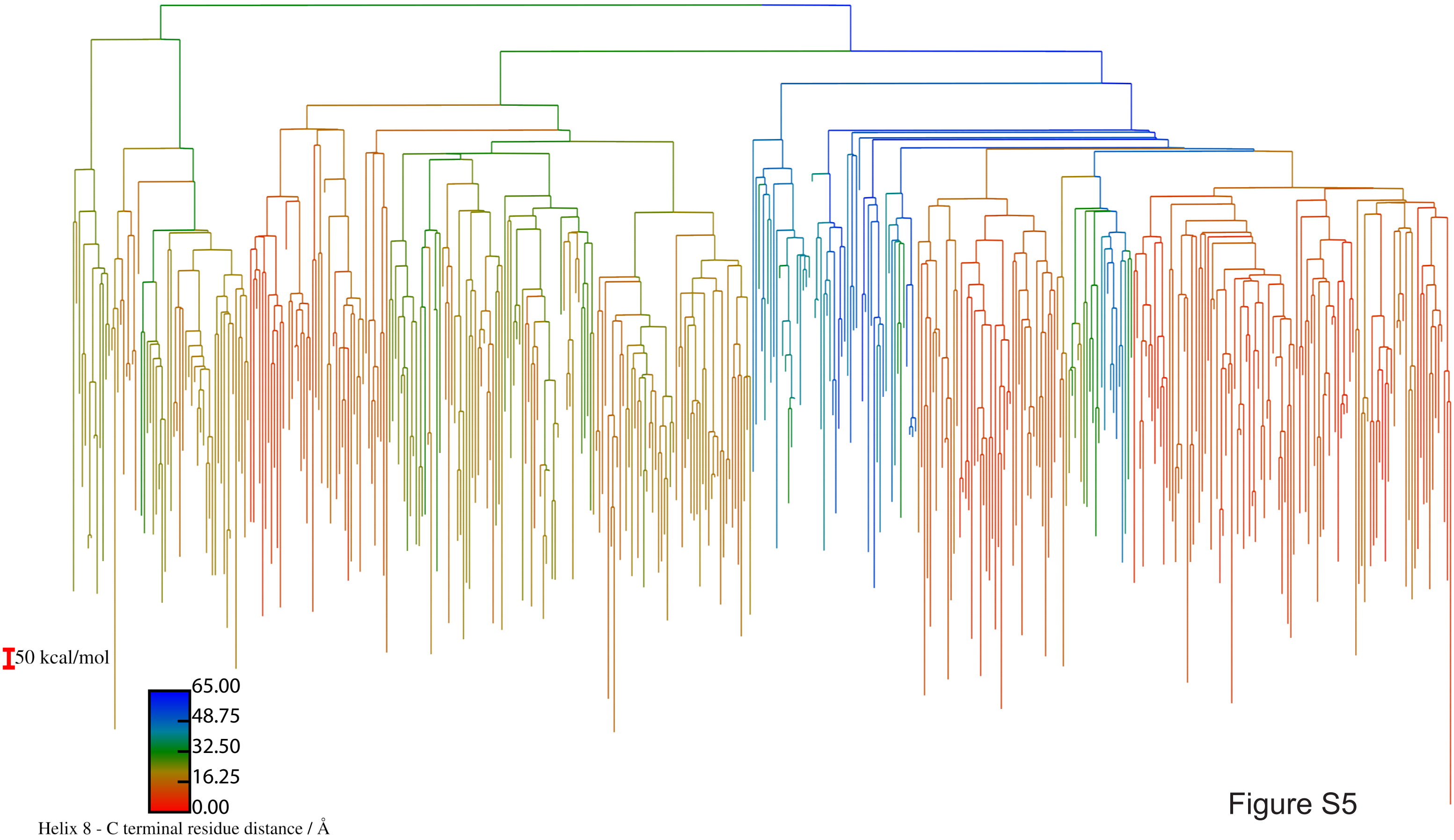

### Supplementary Figure 6

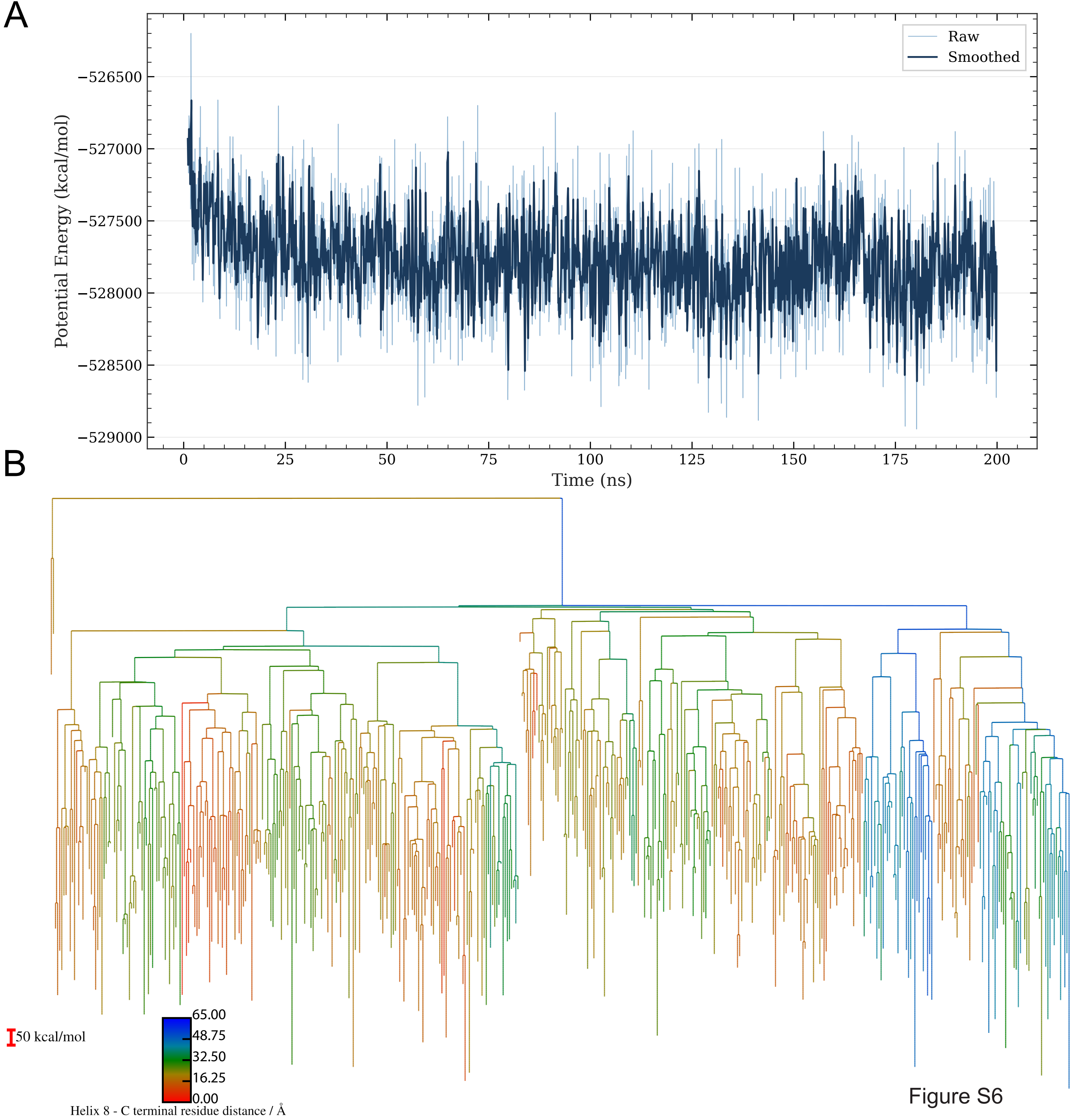

### Supplementary Figure 7

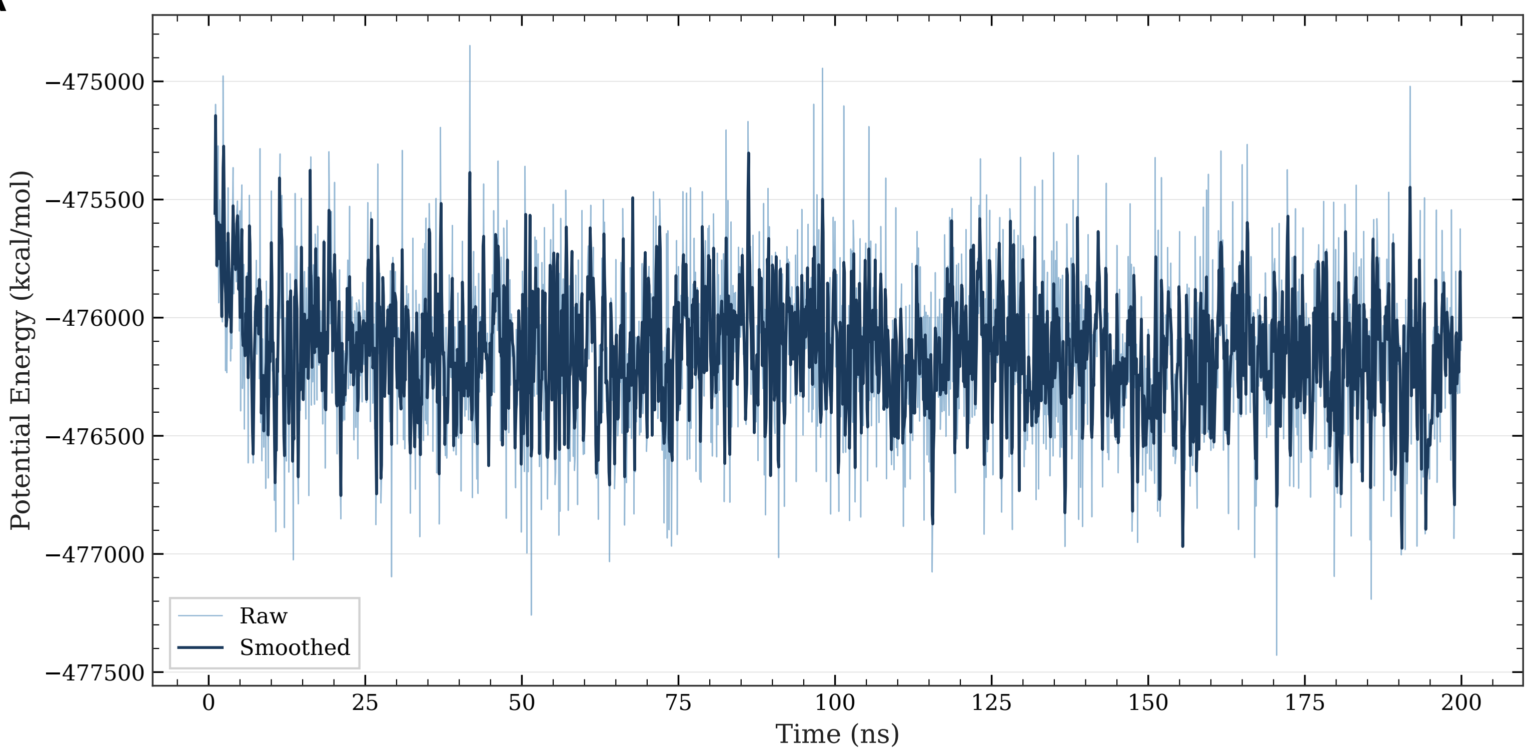

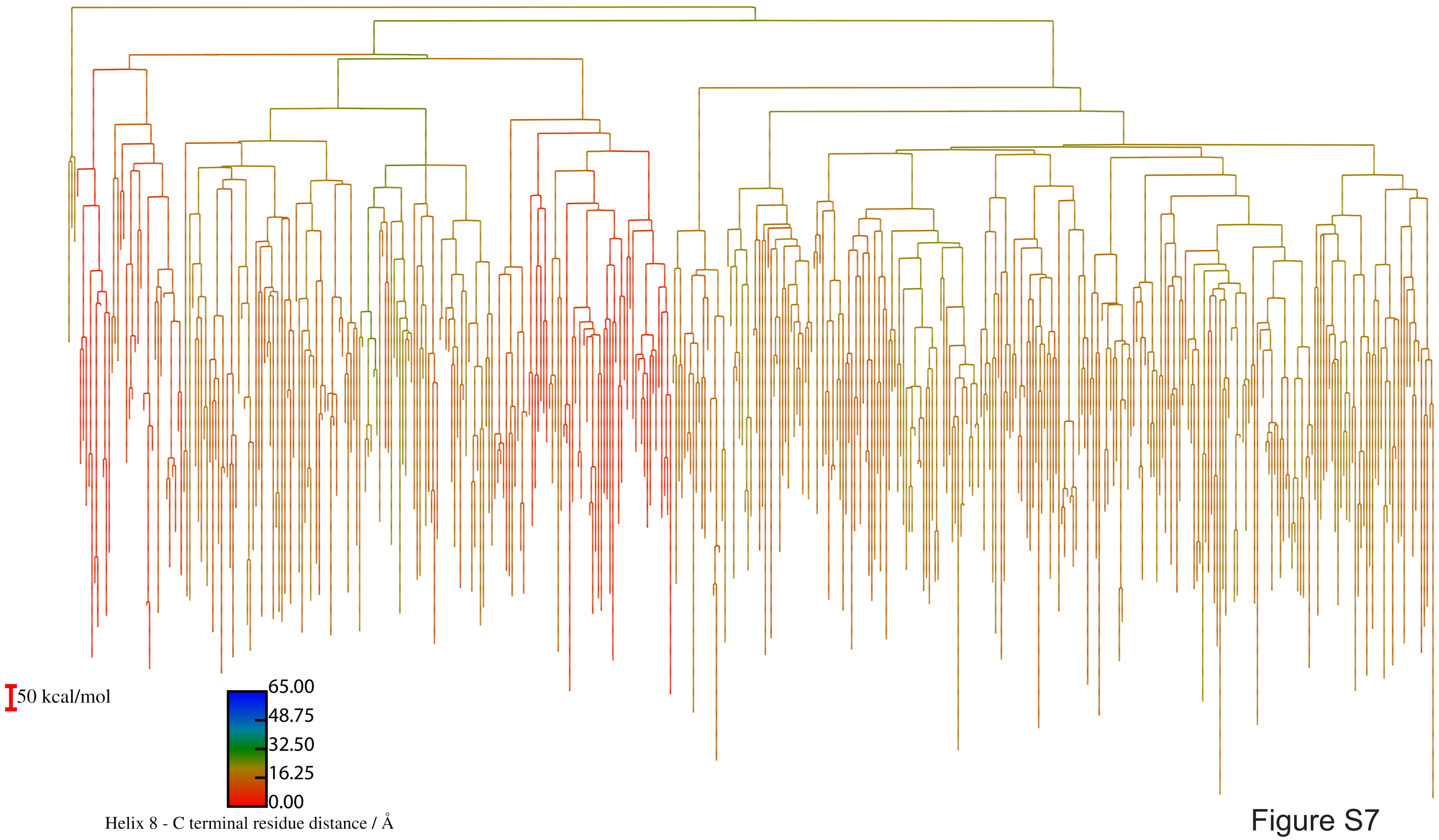

### Supplementary Figure 8

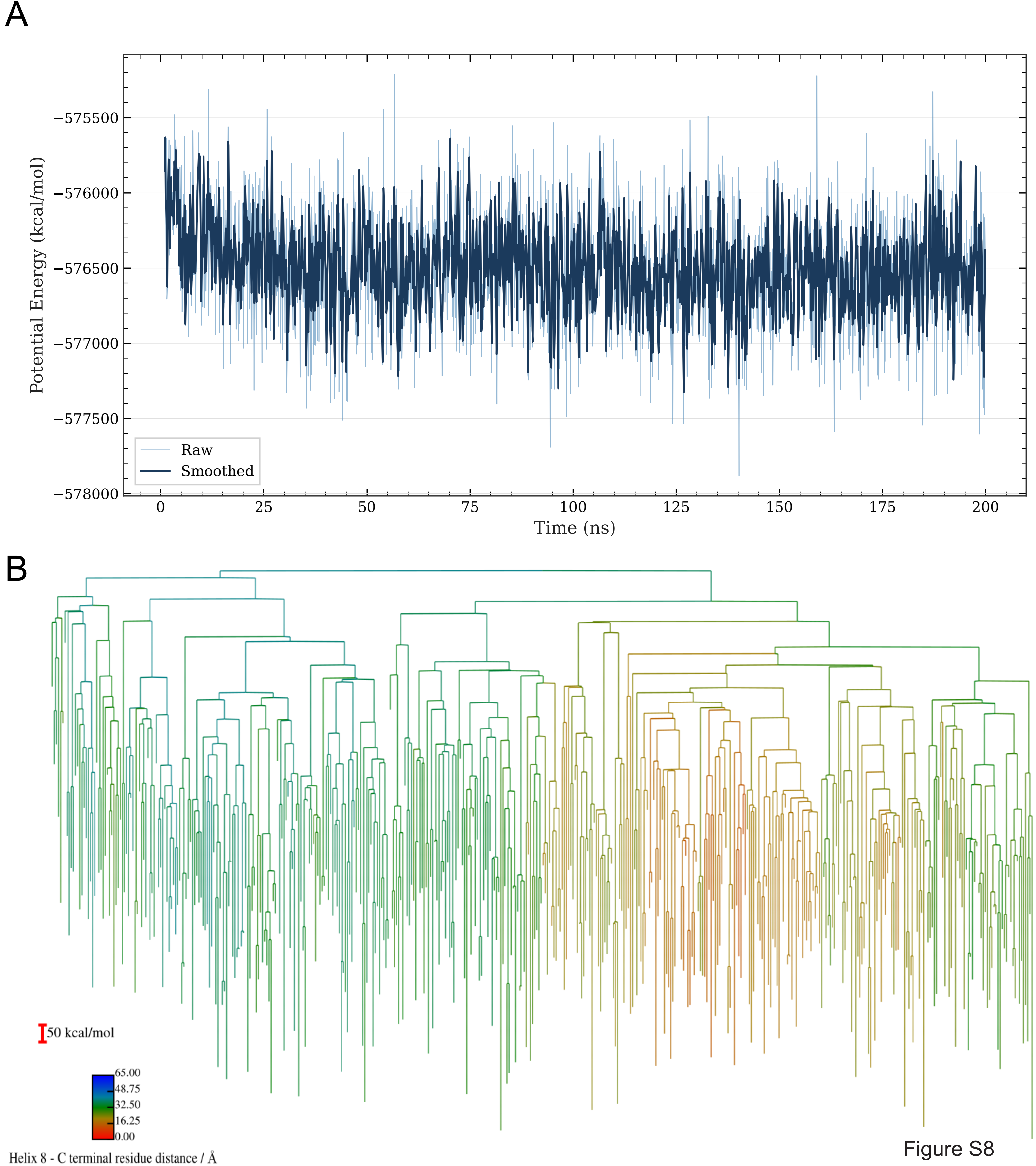

### Supplementary Figure 9

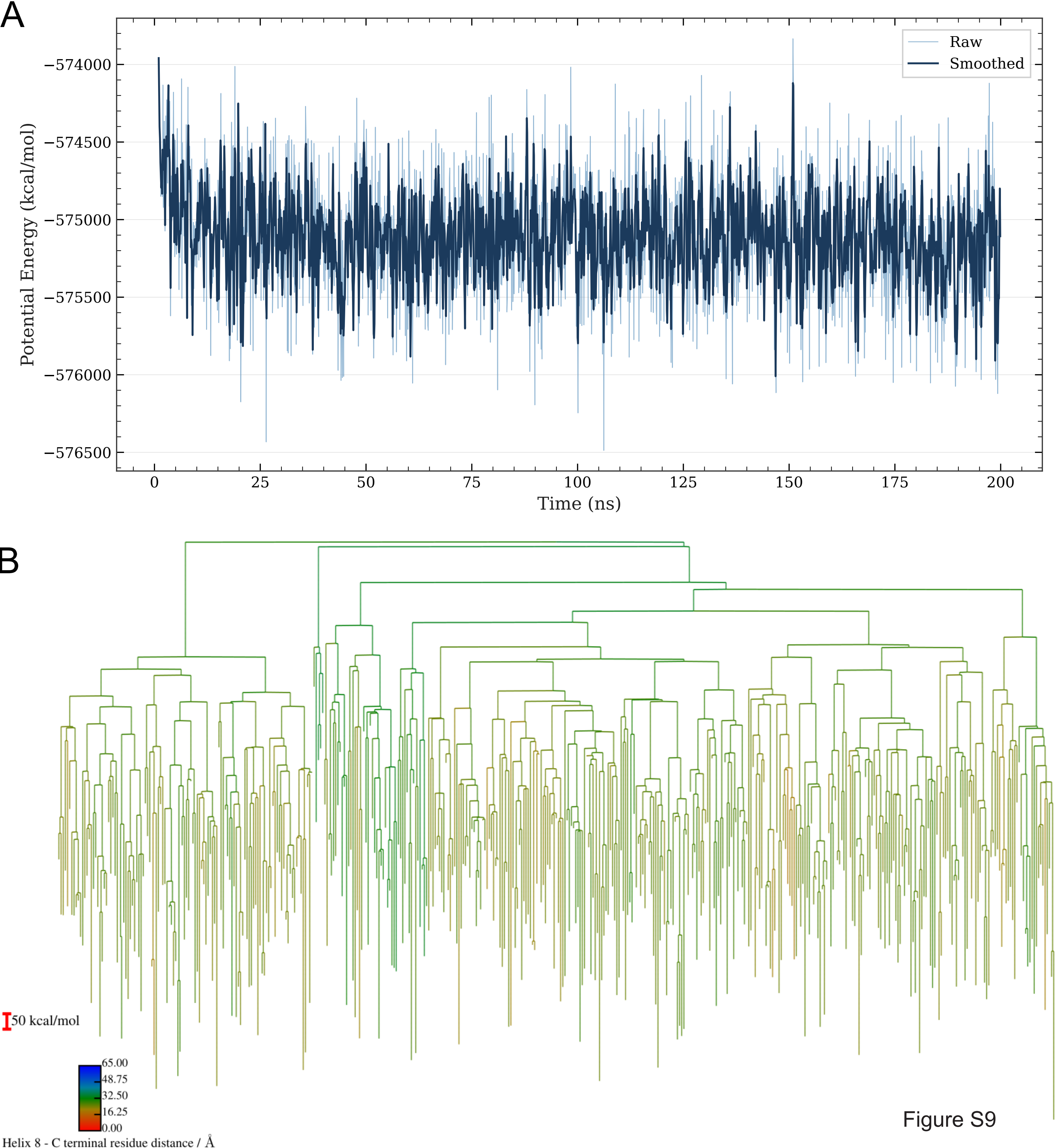
